# Breast cancer stem cell-derived tumors escape from γδ T cell immunosurveillance *in vivo* by modulating γδ T cell ligands

**DOI:** 10.1101/2022.04.04.486931

**Authors:** Katrin Raute, Juliane Strietz, Geoffroy Andrieux, Oliver S. Thomas, Klaus M. Kistner, Marina Zintchenko, Peter Aichele, Houjiang Zhou, Wilfried Weber, Melanie Boerries, Mahima Swamy, Jochen Maurer, Susana Minguet

## Abstract

Triple negative breast cancer (TNBC) lacks targeted therapy options. TNBC is enriched in breast cancer stem cells (BCSCs), which play a key role in metastasis, chemoresistance, relapse and mortality. γδ T cells hold great potential in immunotherapy against cancer, and might be an alternative to target TNBC. γδ T cells are commonly observed to infiltrate solid tumors and have an extensive repertoire of tumor sensing, recognizing stress-induced molecules and phosphoantigens (pAgs) on transformed cells. We show that patient-derived triple negative BCSCs are efficiently recognized and killed by *ex vivo* expanded γδ T cells from healthy donors. Orthotopically xenografted BCSCs, however, were refractory to γδ T cell immunotherapy. Mechanistically, we unraveled concerted differentiation and immune escape: xenografted BCSCs lost stemness, expression of γδ T cell ligands, adhesion molecules and pAgs, thereby evading immune recognition by γδ T cells. Indeed, neither pro-migratory engineered γδ T cells, nor anti-PD-1 checkpoint blockade significantly prolonged overall survival of tumor-bearing mice. BCSC immune escape was independent of the immune pressure exerted by the γδ T cells, and could be pharmacologically reverted by Zoledronate or IFN-α treatment. These results pave the way for novel combinatorial immunotherapies for TNBC.

## Introduction

In breast cancer, the most commonly diagnosed cancer worldwide in 2020 (Sung et al. 2021), breast cancer stem cells (BCSCs) play a key role in metastasis formation, tumor recurrence and mortality of patients (Baccelli et al. 2013; Creighton et al. 2009). Specifically targeting BCSCs is a promising avenue for cancer therapy, yet it faces multiple challenges mainly due to BCSC intrinsic cell heterogeneity and drug resistance (Scioli et al. 2019). BCSC-focused therapies are of even greater importance in the fight against triple negative breast cancer (TNBC), the most aggressive and lethal breast cancer subtype, for which no targeted treatment options are currently available (Bianchini et al. 2016).

Immunotherapies have recently been employed for treating a variety of cancers by transferring autologous *ex vivo*-engineered αβ T cells back into patients. However, the efficacy of these treatments relies on the presence of tumor-specific antigens presented on MHC I molecules. The loss of MHC I expression and the concomitant lack of peptide presentation in cancer cells can undermine conventional αβ T cell recognition, and thus lead to tumor immune escape, which has been widely observed in cancers, including TNBC (Pedersen et al. 2017; Dusenbery et al. 2021; Dhatchinamoorthy et al. 2021). To overcome this limitation, novel immunotherapies using unconventional non-MHC-restricted lymphocytes, such as γδ T cells are currently being investigated. γδ T cells react to stress-induced proteins or phosphoantigens (pAgs) which accumulate in tumor cells due to their deregulated metabolism (Groh et al. 1999; Bathaie et al. 2017; Morita et al. 2007).

Two sub-populations of γδ T cells are currently in focus for therapeutic applications: Vγ9Vδ2 T cell receptor (TCR)-expressing γδ T cells (Vδ2^+^ T cells) and Vδ1^+^ γδ T cells. Vδ2^+^ T cells represent the predominant γδ T cell subset in the human blood and specifically recognize pAgs (Gober et al. 2003; Yoshimasa Tanaka et al. 1995; Constant et al. 1994). In contrast, Vδ1^+^ T cells represent a minor population in peripheral human blood and mainly locate to epithelial tissues. Vδ1^+^ T cells react to several lipid and protein antigens (Groh et al. 1998; Roy et al. 2016; Luoma et al. 2013). Both of these γδ T cell subsets have been described to kill a variety of cancer cell lines upon activation of their specific TCRs, innate receptors like NKG2D, or by engaging the “death receptors” Fas or TRAIL on tumor cells (Sebestyen et al. 2019; Morrow et al. 2019; Spada et al. 2000; D’Asaro et al. 2010). We investigated here the potential of these two γδ T cell subsets to target BCSCs.

Reactivity of γδ T cells against cancer stem cells of several cancer entities has been described (Todaro et al. 2009; Lai et al. 2012; Nishio et al. 2012). Only two recent studies addressed the reactivity of γδ T cells against BCSCs (Chen et al. 2017; Dutta et al. 2021). While Chen et al. did not observe a difference in γδ T cell-mediated cytotoxicity between BCSCs and their non-stem cell counterparts, Dutta et al. reported that BCSCs were less susceptible to being killed by γδ T cells. To reconcile these discrepancies and to shed light onto the potential of γδ T cells to target BCSCs, studies better reflecting the clinical reality are urgently needed.

The potential success of γδ T cell immunotherapy against solid tumors also relies on the efficient localization of these cells into the tumor tissue, since T cells require direct cell-cell contact to exert their cytotoxicity. γδ T cells need to extravasate from the blood stream into the tissue and migrate in the stromal tumor compartment. It has been shown for conventional αβ T cells that the extracellular matrix (ECM), the major non-cellular fraction of the tumor microenvironment, negatively affects the migration and infiltration of αβ T cells in non-small-cell lung cancer and ovarian cancer (Salmon et al. 2012; Bougherara et al. 2015). Indeed, high stromal content, as well as low numbers of infiltrating T cells, have been associated with poor clinical outcome in breast cancer (Vangangelt et al. 2018). Therefore, increasing the infiltration of T cells into tumors is a major goal in immunotherapies. In line with this, exogenous expression of the ECM-modifying enzyme heparanase can increase the tumor infiltration of chimeric antigen receptor (CAR)-expressing αβ T cells, promoting tumor rejection in melanoma and neuroblastoma xenograft models (Caruana et al. 2015). Whether the ECM-rich stromal compartment of tumors also hampers γδ T cell migration, infiltration and tumor rejection has not yet been investigated.

We sought here to use a model aiming to reflect the situation in TNBC human patients to test the effect of γδ T cell-based therapy.

## Results

### Expanded γδ T cells efficiently kill patient-derived triple negative BCSCs

To test the cytotoxic potential of human γδ T cells towards patient-derived triple negative BCSCs, we expanded γδ T cells from peripheral blood mononuclear cells of healthy donors using concanavalin A (ConA) stimulation. As previously described (Siegers et al. 2013), expansion resulted in a specific enrichment of effector memory Vδ1^+^ and Vδ2^+^ T cells (Extended Data Fig. 1a). The expression of the activating natural killer cell receptor NKG2D, which mediates the cytolytic activity of γδ T cells (Wrobel et al. 2007; Lança et al. 2010), was upregulated during γδ T cell expansion (Extended Data Fig. 1b). We observed significantly higher NKG2D expression in Vδ2^+^ compared to Vδ1^+^ T cells during expansion; however, NKG2D expression was comparable at the end of the culture period (28 days). It has been reported that the overt expansion of αβT cells, their genetic modification or their stimulation after transfer to the patient can result in T cell exhaustion and functional failure (Xia et al. 2019; Jiang et al. 2015). Therefore, we followed the expression of the inhibitory receptors PD-1, TIM-3 and LAG-3 throughout the expansion (Extended Data Fig. 1b). PD-1 was upregulated within the first 10 days after ConA stimulation and then declined to basal level. TIM-3, in contrast, exhibited a constant increase in expression as the expansion progressed. LAG-3 expression was drastically increased in the first 10 days after ConA stimulation, was then reduced but remained upregulated until the end of the expansion. The three inhibitory receptors were expressed significantly higher in Vδ2^+^ compared to Vδ1^+^ T cells upon stimulation, and TIM-3 and LAG-3 remained highly expressed in Vδ2^+^ T cells at the end of the observed expansion period. Of note, the percentage of Vδ1^+^ and Vδ2^+^ T cells strongly varied between healthy donors. In the majority of donors, Vδ2^+^ T cells were highly enriched (Extended Data Fig. 1c). Expanded γδ T cells were used for experiments between day 14 and day 35 after starting the expansion since the cells have reached a stable phenotype at this time window.

Next, we determined whether these γδ T cells could kill patient-derived triple negative BCSCs *in vitro*. Expanded γδ T cells from three healthy donors killed all tested patient-derived triple negative BCSC lines (BCSC1, BCSC3 and BCSC5) in an effector to target ratio-dependent manner (Fig. 1a). BCSC5 was most efficiently killed. To test whether the cytotoxicity was mediated by Vδ1^+^ or Vδ2^+^ T cells, we separated these two T cell subsets after expansion (Extended Data Fig. 1d). Both Vδ1^+^ and Vδ2^+^ T cells displayed similar cytotoxicity in response to BCSC1, BCSC3 and BCSC5 (Fig. 1b). Remarkably, BCSCs only induced degranulation in Vδ2^+^ T cells, measured indirectly via CD107a (Fig. 1c). This was most prominent for BCSC5, which led to significantly higher degranulation in Vδ2^+^ T cells compared to BCSC1 and BCSC3. That Vδ1^+^ T cells did not degranulate after BCSC contact was not a consequence of a general incapability of Vδ1^+^ T cells to degranulate since stimulation with α-CD3 and α-CD28 antibodies led to CD107a accumulation in both T cell subsets (Fig. 1c). These results suggest that Vδ1^+^ and Vδ2^+^ T cells kill patient-derived human BCSCs by using different mechanisms.

**Fig. 1:**
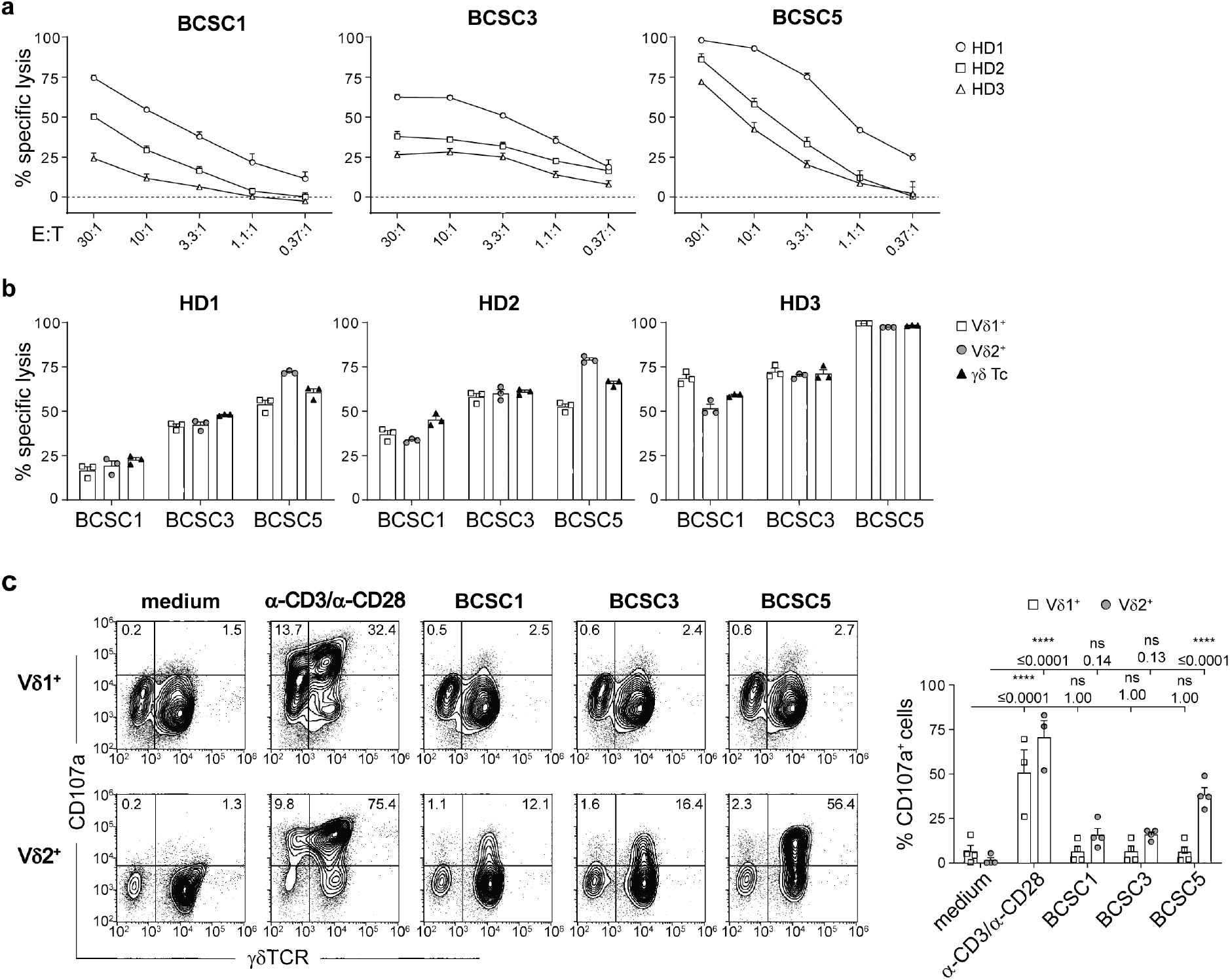
Expanded γδ T cells recognize and kill BCSCs *in vitro*. **(a)** *In vitro* killing of luciferase-expressing BCSCs by γδ T cells after 8 h at various effector to target (E:T) ratios. Results from two independent experiments with a total of three healthy donors (HD) of γδ T cells are shown (means ± SEM). **(b)** *In vitro* killing of BCSCs by γδ T cells and magnetic activated cell sorting (MACS)-separated Vδ1^+^ and Vδ2^+^T cells performed as in (a). Cells were co-cultured at an E:T ratio of 10:1. Results from three healthy donors of γδ T cells are shown (means ± SEM). **(c)** Flow cytometry-based analysis of degranulation by MACS-separated Vδ1^+^ or Vδ2^+^T cells in response to BCSC contact for 3 h. Stimulation with α-CD3 and α-CD28 served as positive control. Representative dot plots (left) and statistical analysis (right) of the percentage of CD107a^+^Vδ1^+^ or CD107a^+^Vδ2^+^cells. Results from two healthy donors of γδ T cells obtained in three independent experiments were pooled (means ± SEM). Two-way ANOVA followed by Sidak’s post hoc test comparing stimulated or co-cultured cells to the corresponding medium control. **** p≤0.0001. E:T, effector to target; HD, healthy donor.

### Expanded γδ T cells are attracted by BCSC-conditioned medium

In addition to recognizing and killing BCSCs, *in vivo* γδ T cells need to be attracted and migrate towards the tumor sites. Therefore, we next assessed the migratory capacity of γδ T cells towards BCSC-conditioned medium in transwell experiments. Conditioned medium from BCSC1, BCSC3 and BCSC5 efficiently attracted γδ T cells (Fig. 2a). The migration of Vδ1^+^ T cells towards BCSC-conditioned medium was increased compared to control medium, but strongly dependent on the T cell donor. In contrast, Vδ2^+^ T cells from all tested healthy donors migrated robustly in response to BCSC-conditioned medium (Fig. 2b). γδ T cells outperformed αβ T cells regarding their migration ability towards BCSC5-conditioned medium (Fig. 2c).

**Fig. 2:**
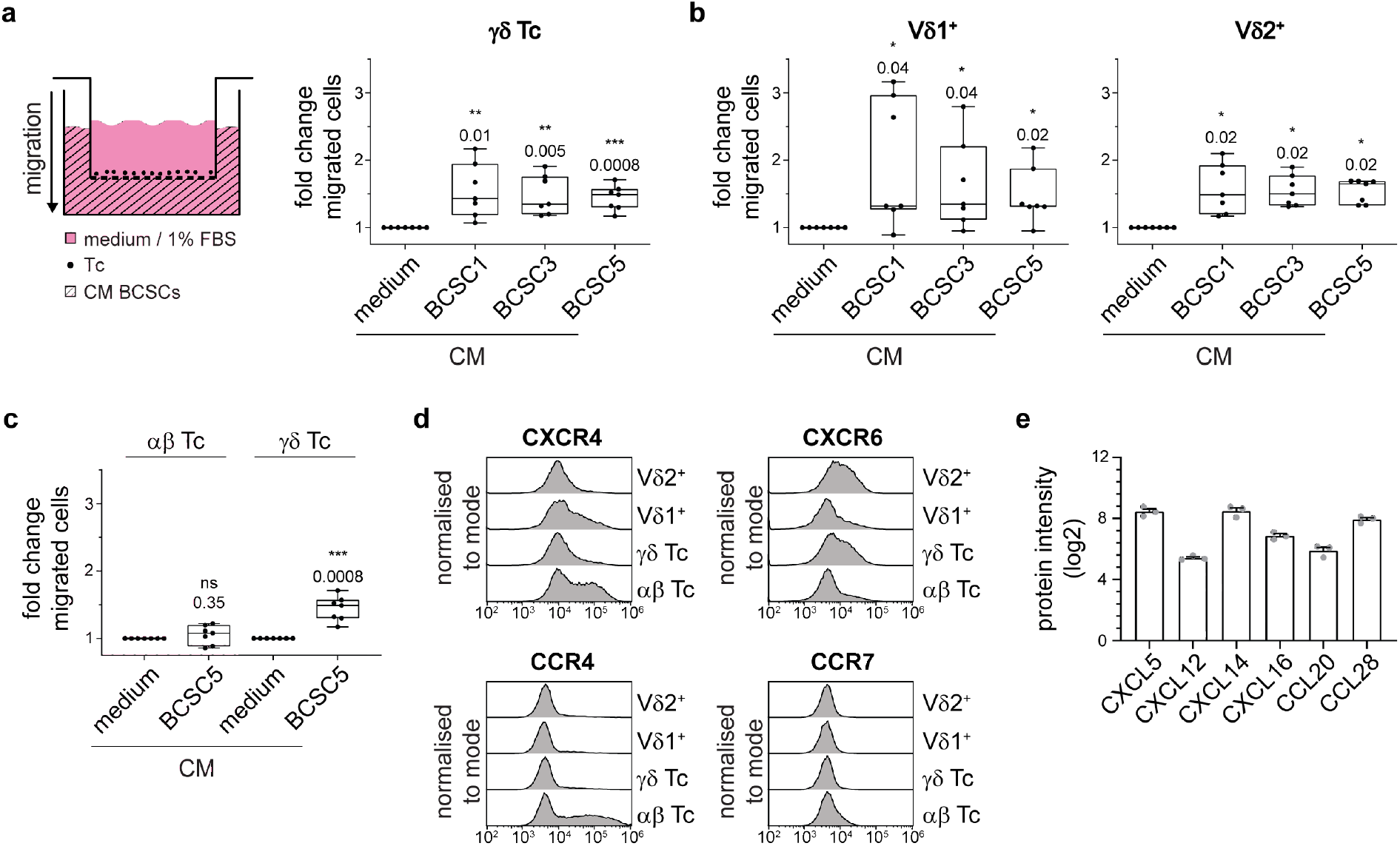
Expanded γδ T cells migrate towards BCSC-conditioned medium. **(a)** Schematic transwell migration assay (left). Primary T cells were seeded in the wells of a permeable support with 5 µm pore size. The lower compartment was filled with BCSC-conditioned medium (CM) and cells that transmigrated into the lower compartment were analyzed by flow cytometry. Statistical analysis of transmigrated γδ T cells (CD3^+^γδTCR^+^) is shown (right). Basal migration towards medium was set to 1.0 and fold changes from two independent experiments using seven healthy donors in total were pooled (median, min to max). One sample t-test against hypothetical value of 1.0. **(b)** Transwell assays were performed and analyzed as in (a). Transmigrated Vδ1^+^ (CD3^+^γδTCR^+^Vδ1^+^) and Vδ2^+^ (CD3^+^γδTCR^+^Vδ2^+^) T cells were distinguished by flow cytometry. One-sample t-test (for Vδ1^+^) or one sample Wilcoxon test (for Vδ2^+^). **(c)** Transwell assays were performed and analyzed as in (a). Comparison of transmigrated αβ T cells (CD3^+^γδTCR^-^) and γδ T cells (CD3^+^γδTCR^+^; data from (a)) in response to BCSC5-conditioned medium. One-way ANOVA followed by Dunnett’s post hoc test comparing BCSC5 CM groups to control medium groups. **(d)** Representative chemokine receptor expression levels in αβ (CD3^+^γδTCR^-^), γδ (CD3^+^γδTCR^+^), Vδ1^+^ (CD3^+^γδTCR^+^Vδ1^+^) and Vδ2^+^ (CD3^+^γδTCR^+^Vδ2^+^) T cells. Shown are representative flow cytometry histograms from one healthy donor out of three healthy donors analyzed. **(e)** Chemokine expression levels determined by quantitative mass spectrometry of BCSC5. Protein intensities (log2) of three replicates are shown (means ± SEM). * p≤0.05, ** p≤0.01, *** p≤0.001.

The different migratory responses exhibited by the specific T cell subsets were more thoroughly investigated. We thus analyzed the expression levels of 12 chemokine receptors (CXCR1, CXCR3, CXCR4, CXCR5, CXCR6, CCR2, CCR3, CCR4, CCR5, CCR6, CCR7 and CCR10). Among those, we found four of them, namely CXCR4, CXCR6, CCR4 and CCR7, to be differentially expressed in αβ, γδ, Vδ1^+^ and Vδ2^+^ T cells (Fig. 2d). The expression of CXCR6 correlated well with the migratory responses observed. The only known ligand for CXCR6 is CXCL16, and CXCL16 was indeed detected in a proteomic approach using BCSC5 culture cells (Fig. 2e). Besides CXCL16, we identified the chemokines CXCL12, CXCL5, CXCL14, CCL20 and CCL28 to be expressed by BCSC5 (Fig. 2e). However, the chemokine expression pattern alone might be insufficient to identify the chemokine-chemokine receptor pairs involved in the attraction of T cells by BCSC-conditioned medium due to the high degree of promiscuity defining the chemokine system. For example, even the migration towards the well-known T cell attractant CXCL12 only partially correlated with the expression levels of its best-characterized receptor CXCR4 (Extended Data Fig. 2a and b).

Taken together, ConA-expanded γδ T cells most efficiently recognize and kill BCSC5 among the tested BCSCs, and exhibit a robust migration towards BCSC5-conditioned medium. Therefore, we focused our studies on BCSC5 and conducted further experiments with γδ T cell cultures containing mainly Vδ2^+^ T cells and less than 10% of Vδ1^+^ T cells to minimize heterogeneity.

### MMP14 expression in γδ T cells increases their migration capacity in ECM-rich environments

Our transwell assays demonstrated that γδ T cells are efficiently attracted towards BCSC5-conditioned medium. However, BCSC-derived xenotransplanted tumors are surrounded by a dense ECM, similar to the original TNBC tumors (Metzger et al. 2017). This ECM-rich stromal compartment of tumors might hamper γδ T cell migration and infiltration. Therefore, we aimed to boost γδ T cell migration by expressing the membrane-anchored matrix metalloprotease 14 (MMP14) in γδ T cells. MMP14 is one of the 26 known endopeptidases of the human MMP protein family (Kapoor et al. 2016). It can cleave a plethora of ECM proteins like fibronectin, collagen (type I, II and III) and laminin (Pei and Weiss 1996; Ohuchi et al. 1997; Koshikawa et al. 2005; d’Ortho et al. 1997). Furthermore, MMP14 has been intensively studied in the process of tumor cell migration where it was shown to possess pro-migratory functions (Hotary et al. 2000; Sabeh et al. 2004). MMP14 is indeed endogenously expressed in γδ T cells directly after ConA stimulation but is downregulated over time. In contrast, αβ T cells did not upregulate MMP-14 upon activation (Extended Data Fig. 3a). To maintain MMP14 expression, expanded γδ T cells were lentivirally transduced with a mock vector, MMP14 or the catalytically inactive mutant MMP14^E240A^, and expression was verified by flow cytometry (Fig. 3a, Extended Data Fig. 3b). We then assessed the effect of MMP14 on γδ T cell migration towards CXCL12 and FBS in Matrigel, a model matrix for basement membranes (Fig. 3b). MMP14 expression increased the percentage of γδ T cells migrating further than 10 or 100 µm, while the catalytically inactive mutant failed to promote migration (Fig. 3b). These findings demonstrate that MMP14 promotes the migration of γδ T cells in basement membrane-like ECM.

**Fig. 3:**
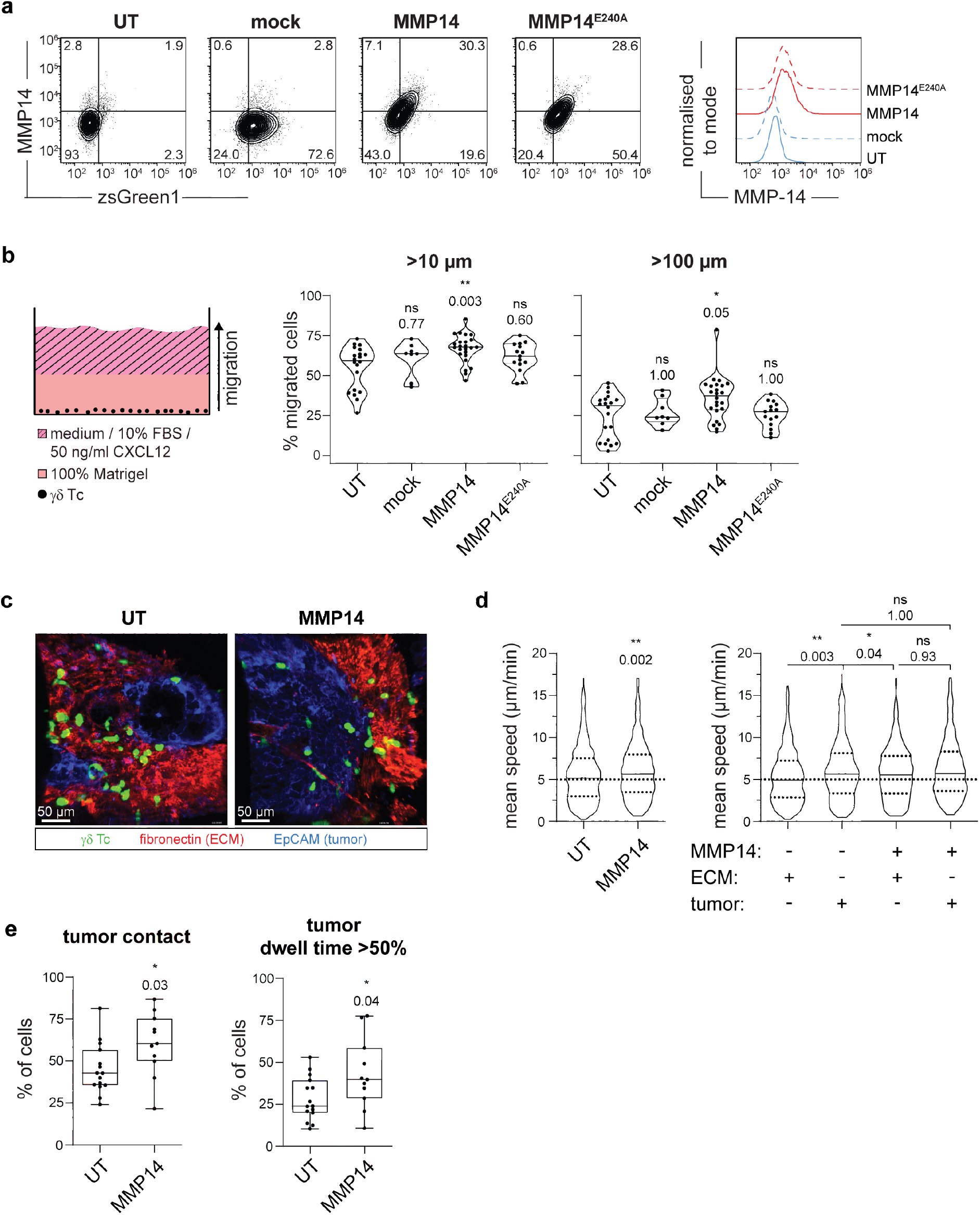
MMP14 expression increases γδ T cell migration in basement membrane-like Matrigel and in BCSC5 tumor tissue. **(a)** Representative dot plots (left) and histograms (right) of γδ T cells expressing mock, MMP14 or MMP14^E240A^ on day 3 after transduction with a multiplicity of infection (MOI) of 5. Untransduced (UT) cells served as control. **(b)** Schematic illustration of the 3D migration assay in Matrigel (left). γδ T cells were seeded into ibidi µ-angiogenesis slides in 100% Matrigel. Migration towards medium supplemented with 10% FBS and 50 ng/ml CXCL12 was assessed via confocal microscopy after 48 h. Statistical analysis (right) of the percent of γδ T cells migrating further than 10 and 100 µm (medians are indicated). Results of 8-24 wells per condition from five independent experiments are shown. Kruskal-Wallis test followed by Dunn’s post hoc test comparing transduced to untransduced cells. **(c)** Migration of CMFDA-labeled UT (see also Supplementary Video 1) or MMP14 expressing (see also Supplementary Video 2) γδ T cells (green) in vibratome sections of viable BCSC5 xenograft tumors. Shown are representative Z-projections of confocal time-lapse videos from BCSC5 xenograft tumor slices stained for EpCAM (blue) and fibronectin (red) to identify tumor cell regions and stromal compartments, respectively. **(d)** The mean speed of γδ T cells migrating in BCSC5 xenograft tumor slices is shown (left; Mann-Whitney test). The same data were analyzed with respect to the mean speed of γδ T cells in stromal ECM compartments (fibronectin^+^) and tumor cell regions (EpCAM^+^) of BCSC5 xenograft tumor slices (right; Kruskal-Wallis test followed by Dunn’s post hoc test comparing all groups against each other). **(e)** The percentage of cells, which have been able to enter EpCAM^+^ tumor tissue (left) or which have resided over 50% of their monitored time inside EpCAM^+^ tumor tissue (right), was quantified in each time-lapse experiment (median, min to max). Unpaired t-test, two-tailed. (D and E) Results from 15 (UT) and 11 (MMP14) time-lapse acquisitions from seven independent experiments are shown including at least 440 tracks per group. Outliers were identified and removed using the ROUT method (Q=2%). * p≤0.05, ** p≤0.01, *** p≤0.001.ECM, extracellular matrix; EpCAM, epithelial cellular adhesion molecule; UT, untransduced.

To test whether the pro-migratory function of MMP14 observed in Matrigel also supports interstitial migration in tumors, we analyzed γδ T cell migration in viable slices of BCSC5 xenograft tumors via confocal microscopy following published protocols (Extended Data Fig. 3c) (Salmon et al. 2012; Salmon et al. 2011). For this purpose, BCSC5 cells were orthotopically transplanted into the mammary fat pad of NOD SCID mice as previously described (Li et al. 2020; Metzger et al. 2017; Strietz et al. 2021). CMFDA-labeled non-transduced γδ T cells or γδ T cells expressing MMP14 were plated on top of unfixed xenograft-derived tumor slices and were microscopically monitored. EpCAM and fibronectin staining served to distinguish tumor islets and ECM-rich stroma, respectively (Fig. 3c and Supplementary Video 1 and 2). MMP14 expression increased the interstitial migration speed of γδ T cells when compared to non-transduced cells (Fig. 3d, left). When differentially analyzing the migration speed of γδ T cells within the tumor tissue or the stromal ECM, we observed that non-transduced γδ T cells migrated faster in the tumor tissue than in the stromal ECM (Fig. 3d). This is in line with observations made for αβ T cells in ovarian and lung cancer (Peranzoni et al. 2018; Bougherara et al. 2015). MMP14 expression increased the average speed of γδ T cells in the ECM to the speed exhibited in the tumor tissue, supporting the functional role of MMP14 in cleaving the tumor-associated ECM (Fig. 3d, right). In addition, a higher percentage of cells per time-lapse experiment was in direct tumor contact when the cells expressed MMP14 (Fig. 3e) and a bigger fraction of γδ T cells resided in the tumor for more than 50% of their monitored time when expressing MMP-14 (Fig. 3e, right). Altogether, these results show that the protease MMP14 boosts the migration of γδ T cells in the ECM-enriched tumor environment, and might thereby help to overcome limitations that hamper γδ T cell migration *in vivo* and, ultimately, therapy outcome.

### γδ T cells fail to control BCSC5 orthotopic xenografts

Now that we had established a system to maximize γδ T cells targeting of BCSC *ex vivo*, we assessed whether γδ T cells could control BCSC5 tumor growth *in vivo*. To this end, we generated BCSC5 orthotopic mammary gland xenografts in NOD SCID mice as described (Metzger et al. 2017; Strietz et al. 2021; Li et al. 2020). Mice were treated with a vehicle control, γδ T cells or γδ T cells expressing MMP14 once the tumors had reached a volume of at least 4 mm^3^ (Extended Data Fig. 4a). The median survival of the mice after treatment start increased from 28 days for vehicle-treated animals to 33 and 38 days for γδ T cell- or γδ T cell MMP14-treated mice, respectively. Although a clear tendency towards increased survival times of the mice treated with γδ T cells or γδ T cells expressing MMP14 was observed, there was no significant increase in the overall survival between the different treatment groups (Fig. 4a). Furthermore, the treatment did not significantly affect the growth kinetics of the individual tumors, although a higher proportion of tumors slowed down their growth when MMP14 was expressed in γδ T cells (Fig. 4a, right). Importantly, neither treatment with γδ T cells nor with γδ T cells expressing MMP14 increased tumor cell invasion or metastasis when compared to vehicle-treated animals. Similar results were obtained using more immunocompromised mice, namely Rag2^-/-^γc^-/-^ (data not shown).

**Fig. 4:**
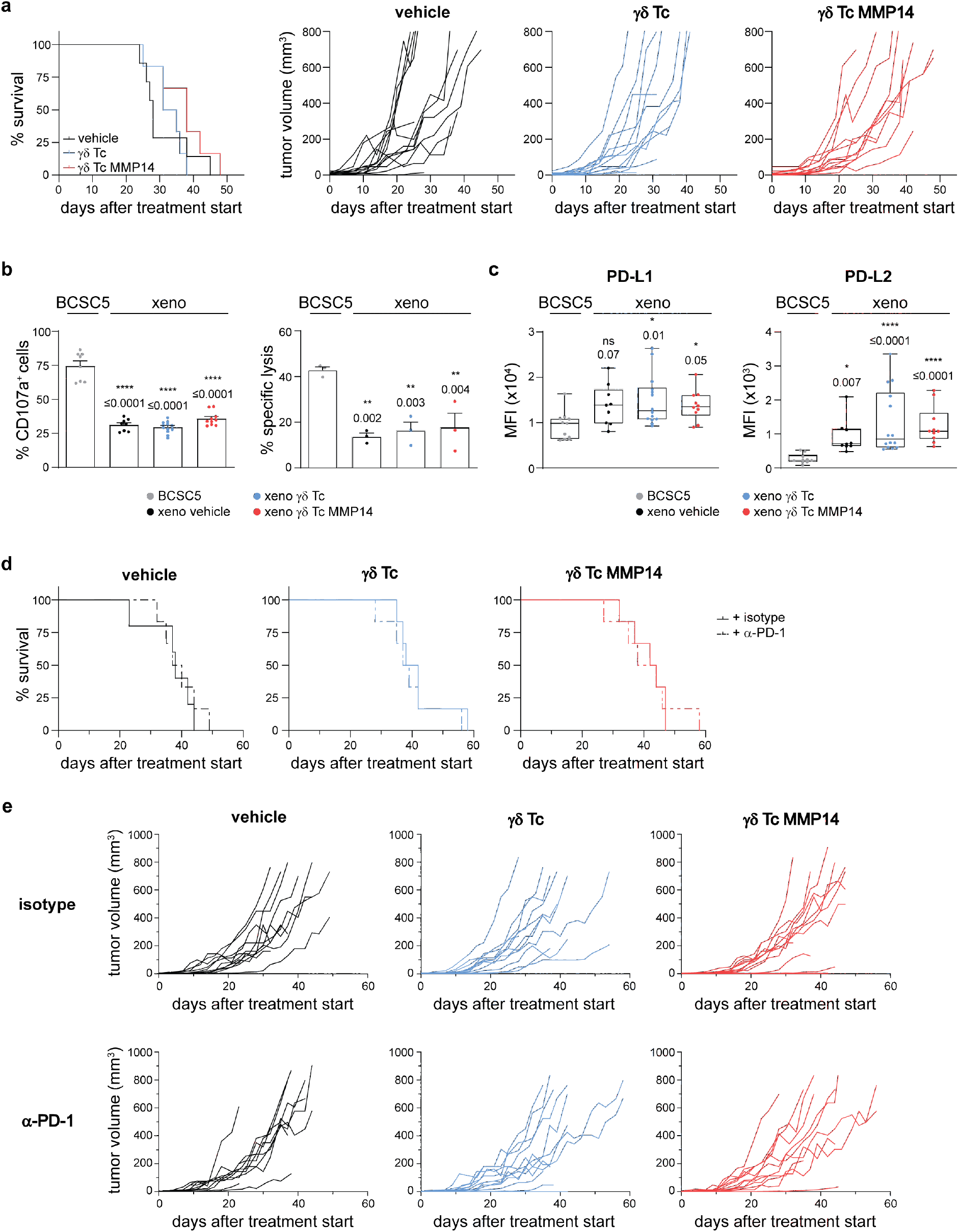
γδ T cells fail to control BCSC5 xenografts in NOD SCID mice. **(a)** Kaplan-Meier plot (left) of BCSC5 xenograft-bearing mice upon treatment with γδ T cells (blue), γδ T cells expressing MMP14 (red) or vehicle control (black) (n=6-7 mice per group). Differences were not statistically significant, Log-rank test (Mantel-Cox). BCSC5 tumor growth curves (right) for individual mice after treatment start. **(b)** γδ T cell-mediated degranulation (left) and cytotoxicity (right) of BCSC5 culture cells or xenograft-derived tumor cells (xeno). Xenograft-bearing mice were treated withγδ T cells, γδ T cells expressing MMP14 or vehicle. For the degranulation assay, γδ T cells were co-cultured with the respective tumor cells for 3 h. The percentage of CD107a^+^ cells of Vδ2^+^-gated cells is shown (means ± SEM). Four to six tumors were analyzed per group and each tumor was tested with two healthy donors of γδ T cells. For i*n vitro* killing of ^51^Cr-labeled tumor cells, γδ T cells were co-cultured with the target cells for 5 h at an E:T ratio of 30:1.Results from three independent experiments with a total of three healthy donors of γδ T cells and three tumors per group were pooled (means ± SEM).One-way ANOVA followed by Dunnett’spost hoc test comparing xenograft to culture cells. **(c)** Flow cytometry-based analysis of PD-L1 and PD-L2 expression levels in xenograft-derived EpCAM^+^ tumor cells (median, min to max). MFIs of 9-14 tumors per group are shown. Kruskal-Wallis test followed by Dunn’s post hoc test comparing xenograft to culture cells. **(d)** Kaplan-Meier plots of NOD SCID mice upon treatment with vehicle control (black), γδ T cells (blue) orγδ T cells expressing MMP14 (red) in combination with an α-PD-1 (Nivolumab) or isotype control antibody (n=5-6 per group). Treatment start was defined for each mouse individually when the first tumor reached a volume of at least 4 mm^3^. 5x10^6^γδ T cells were injected intravenously three times per week. In addition, mice received 0.6x10^6^ IU IL-2 (Proleukin S) on the day of treatment start and every 21 days until the end of the experiment. The end of the experiment was defined by a tumor volume of 800 mm^3^. No significant differences were obtained, Log-rank (Mantel-Cox) test.**(e)** BCSC5 tumor growth curves for individual mice upon treatment with γδ T cells (blue), γδ T cells expressing MMP14 (red) or vehicle control (black) in combination with anα-PD-1 or isotype control antibody (n=5-6 per group).

These results raised the question whether BCSC5 xenograft-derived tumor cells could still be recognized and killed by γδ T cells. Thus, we isolated tumor cells from vehicle-, γδ T cell- or γδ T cell MMP14-treated mice after tumor resection and analyzed the degranulation capacity of γδ T cells in response to these xenograft-derived tumor cells (indicated as “xeno” in the figures). Indeed, the amount of degranulating γδ T cells was significantly reduced from 75% to approximately 30% when compared to the response towards *in vitro* cultured BCSC5 (Fig. 4b, left). Similarly, we observed reduced γδ T cell-mediated killing of xenograft-derived tumor cells compared to BCSCs from the culture (Fig. 4b, right). Importantly, these effects were inherent to the tumor cells and not influenced by the γδ T cell treatment of the tumor-bearing mice (Fig. 4b). We also determined whether secreted soluble molecules decreased the response of γδ T cells to xenograft-derived tumor cells. We analyzed γδ T cell viability and γδ T cell functionality after culturing the cells in conditioned medium of BCSC5 culture cells or xenograft-derived tumor cells for 24 h as previously described (Dutta et al. 2021). γδ T cell viability was not influenced by conditioned medium from either cell type (Extended Data Fig. 4b). Likewise, neither degranulation nor the cytotoxic response of γδ T cells to BCSC5 was affected by the conditioned medium (Extended Data Fig. 4c and d). These findings indicate that the reduced γδ T cell function in response to xenograft-derived tumor cells is not mediated by the secretion of soluble immunosuppressive molecules.

As mentioned before, triggering of inhibitory receptors such as PD1 by their ligands can result in the functional inhibition of T cells. Interestingly, the ligands for the inhibitory receptors PD-1, PD-L1 and PD-L2, were upregulated on xenograft-derived tumor cells (Fig. 4c). To assess whether PD-1 blockade improves γδ T cell treatment *in vivo*, we combined the adoptive transfer of γδ T cells with biweekly applications of a clinically relevant α-PD-1 antibody (Nivolumab). However, the combinatorial treatment of xenograft-bearing mice did not improve the overall survival of the mice but showed a slight reduction in the tumor growth kinetics *in vivo* (Fig. 4d and e). These results suggest that blocking PD-1 is not sufficient to significantly induce γδ T cell-mediated tumor rejection and that other mechanisms might be involved in the immune escape of BCSC5-derived xenografts.

### γδ T cells are efficiently attracted by xenograft-derived tumor cells

To discover proteins and mechanisms involved in the xenograft immune escape *in vivo*, we utilized high-resolution tandem-mass tag-based mass spectrometry to compare the xenograft-derived tumor cell proteomes of all treatment groups with the proteome of BCSC5 culture cells. These analyses revealed that tumor growth in NOD SCID mice drastically changed the proteome of the cells. Around 5600 proteins were differentially expressed by vehicle-treated xenograft cells compared to BCSC5 culture cells (Fig. 5a). Firstly, we more closely examined the chemokine secretion profiles of xenografted BCSC5 compared to culture cells in order to exclude the possibility that xenograft-derived cells lose their ability to attract γδ T cells to the tumor site *in vivo.* Of the detected chemokines, CXCL5 expression was reduced in the xenograft-derived tumor cells compared to BCSC5 culture cells, while CCL20 and CCL28 expression was augmented (Fig. 5b). The rest of the detected chemokines remained unchanged (CXCL12, CXCL14 and CXCL16). None of the observed changes could be associated with a specific treatment. Next, we analyzed γδ T cell migration towards BCSC5 culture cells and freshly isolated xenograft-derived tumor cells. Indeed, cells derived from the xenograft attracted γδ T cells more efficiently than BCSC5 culture cells (Fig. 5c). We assayed Vδ1^+^ and Vδ2^+^ T cells separately and observed that Vδ2^+^ T cells migrated more efficiently towards cells derived from the xenograft compared to Vδ1^+^ T cells (Fig. 5d). These results highlight that xenografts escaped γδ T cell immunotherapy by other means than reducing γδ T cell attraction.

**Fig. 5:**
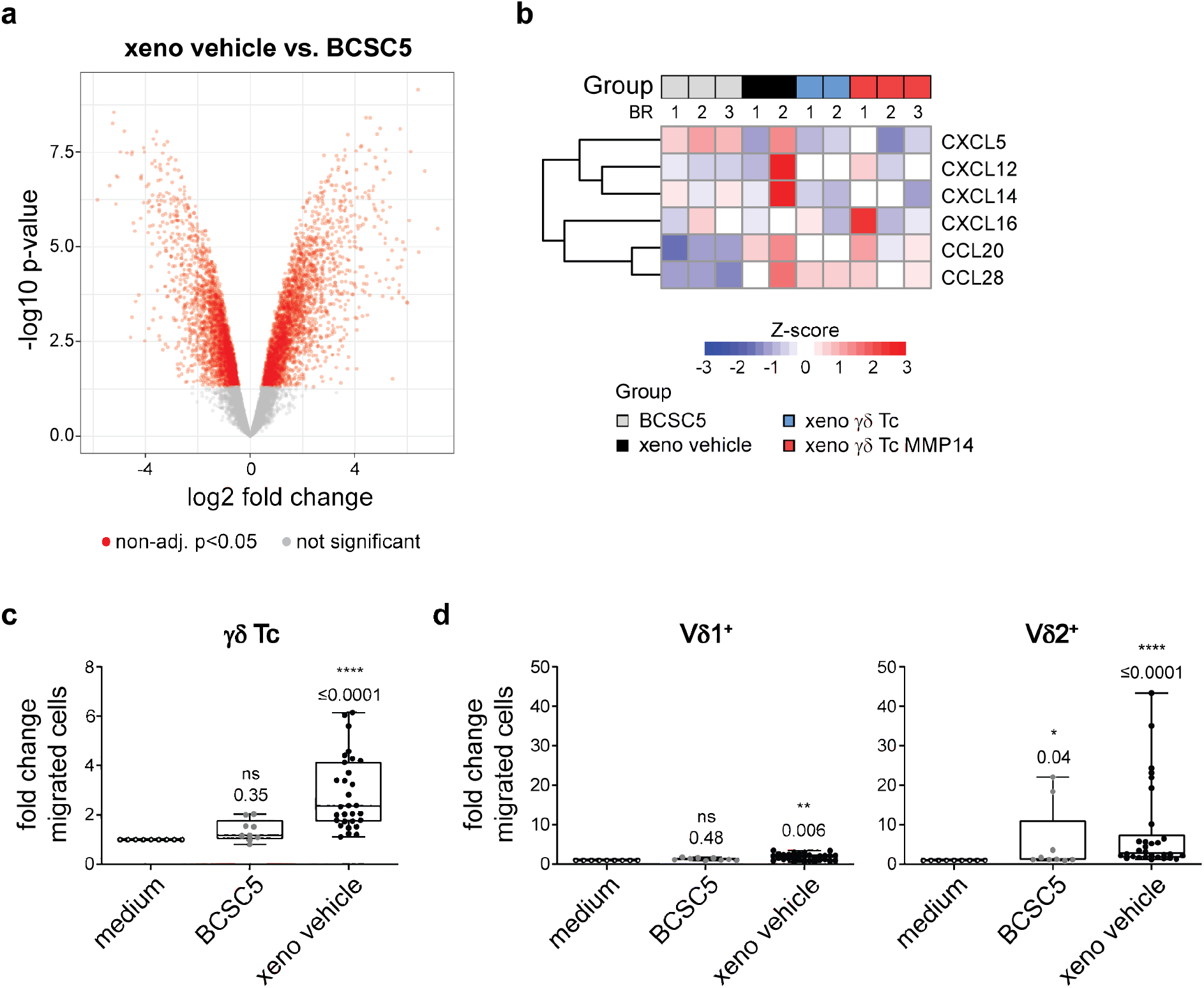
BCSCs phenotypically change *in vivo*, but still induce γδ T cell migration. **(a)** The proteomes of BCSC5 culture cells or freshly isolated xenograft-derived tumor cells were analyzed by mass spectrometry. Volcano plot showing differentially expressed proteins. Proteins regulated with p<0.05 are depicted in red. **(b)** Row-wise Z-score heatmaps showing up- and downregulated proteins in BCSC5 culture cells or xenograft-derived tumors from 2 or 3 biological replicates (BR) as indicated. **(c)** Migration of γδ T cells (CD3^+^γδTCR^+^) in response to BCSC5 culture cells and xenograft derived tumor cells was determined in a transwell assay. Basal migration towards medium was set to 1.0 and fold changes from three independent experiments using the same three healthy donors in each experiment were pooled and analyzed using one-sample Wilcoxon test. **(d)** Migration of Vδ1^+^ (CD3^+^γδTCR^+^Vδ1^+^) and Vδ2^+^(CD3^+^γδTCR^+^Vδ2^+^) T cells in response to BCSC5 culture cells and xenograft derived tumor cells was determined in a transwell assay. Basal migration towards medium was set to 1.0 and fold changes from three independent experiments using the same three healthy donors in each experiment were pooled. One-sample t-test (for Vδ1^+^) or one sample Wilcoxon test (for Vδ2^+^). * p≤0.05, ** p≤0.01, **** p≤0.0001.

### BCSC5 cells differentiate *in vivo* and downregulate the expression of γδ T cell ligands

Next, we performed pathway analyses on our proteomic data and verified that BCSC5 culture cells exhibited a mammary stem cell signature. Remarkably, proteins usually expressed by mammary stem cells were significantly downregulated in xenograft-derived tumor cells and, vice versa, proteins usually lowly expressed in mammary stem cells were upregulated (Fig. 6a). These findings indicate that BCSC5 differentiates *in vivo* losing their breast stem cell signature, despite BCSC-derived preserving the patient’s original molecular tumor subtype (Metzger et al. 2017; Strietz et al. 2021).

**Fig. 6:**
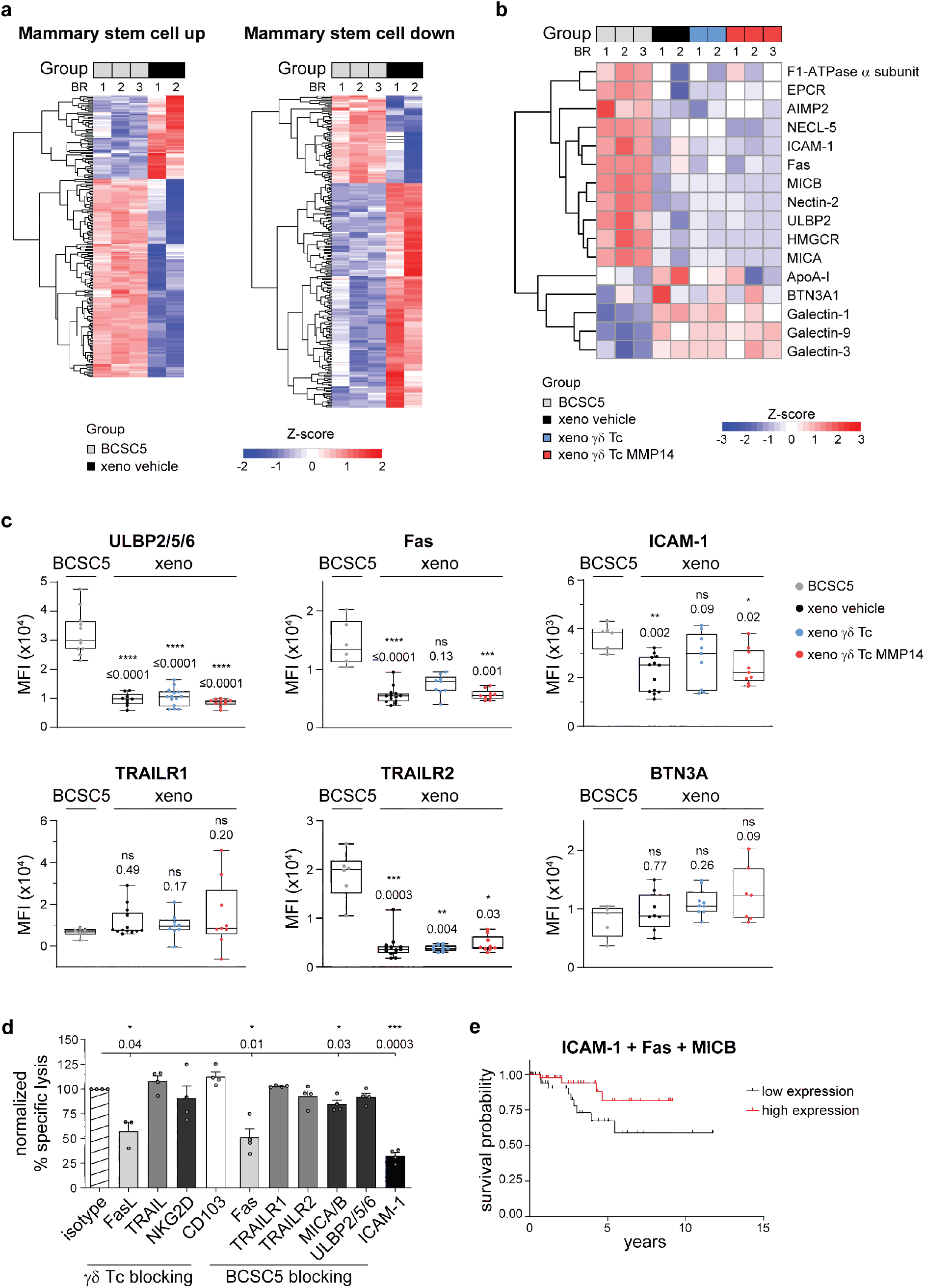
BCSCs differentiate *in vivo* and downregulate the expression of proteins recognized by γδ T cells. **(a)** Row-wise Z-score heatmaps showing up- and downregulated proteins in BCSC5 culture cells or xenograft-derived tumors from 2 or 3 biological replicates as indicated (BR). **(b)** Row wise Z-score heatmap for proteins involved in target cell recognition by γδ T cells or in immune response modulation. **(c)** Flow cytometry-based analysis of BCSC5 culture cells and xenograft-derived tumor cells. MFIs of at least five tumors per group are shown (median, min to max). One-way ANOVA followed by Dunnett’s post hoc test comparing xenograft-derived cells to culture cells (for ULBP2/5/6, ICAM-1 and BTN3A). Kruskal-Wallis test followed by Dunn’s post hoc test comparing xenograft-derived cells to culture cells (for Fas, TRAILR1 and TRAILR2). **(d)** Killing of BCSC5 by γδ T cells requires multiple ligand-receptors interactions. *In vitro* killing of luciferase-expressing BCSC5 cells by γδ T cells. γδ T cells or BCSC5 culture cells were pre-treated with the indicated blocking antibodies for 1 h. Cells were co-cultured at an E:T ratio of 10:1 for 8 h in the presence of the blocking antibodies. Corresponding receptor-ligand pairs are represented in the same color. Isotype antibody control was set to 100% for normalization in each experiment. Results for three to four healthy donors of γδ T cells from three independent experiments were pooled (means ± SEM). One-sample t-test against hypothetical value of 100. **(e)** Cox regression of progression-free survival for TNBC patients sorted by high (upper-quartile) and low (lower-quartile) average clustered expression of the Fas, MICB and ICAM-1. β cox coefficient was - 0.33 and the cox p value was 0.07 suggesting that high expression of the three proteins correlates with a better survival prognosis. * p≤0.05, ** p≤0.01, *** p≤0.001, **** p≤0.0001.

In addition to changes affecting stemness, the transfer and growth of BCSC5 *in vivo* drastically changed the expression levels of well-known γδ T cell ligands and other molecules involved in cancer cell recognition by γδ T cells. We found that a variety of these proteins were specifically downregulated in xenograft-derived tumor cells and that these changes were independent of the γδ T cell-treatment (Fig. 6b). We validated the reduced expression of ULBP-2/5/6, Fas and ICAM-1 via flow cytometry (Fig. 6c). In contrast, MICA/B surface levels were unchanged in the differentiated xenograft cells (data not shown). All these proteins have been previously described to be involved in tumor cell killing by γδ T cells (Uchida et al. 2007; Liu et al. 2009; Rincon-Orozco et al. 2005; Bryant et al. 2011; Li et al. 2011; Spada et al. 2000). Although TRAILR1 and TRAILR2 were not identified in our proteomic study, TRAIL-TRAILR interactions induce γδ T cell-mediated killing (Dokouhaki et al. 2013; D’Asaro et al. 2010). While TRAILR1 expression was not significantly changed on xenograft-derived tumor cells compared to BCSC5 culture cells, TRAILR2 levels were drastically reduced (Fig. 6c). Taken together, the loss of the above-mentioned proteins might explain why γδ T cells cannot efficiently recognize and kill xenograft-derived tumor cells. We next analyzed “The Cancer Genome Atlas” (TCGA; https://www.cancer.gov/tcga) datasets for TNBC patients scoring for the expression level of Fas, MICA/B, TRAILR1/2 and ICAM-1 individually. Cox proportional hazards analysis failed to reveal significant prolonged survival for patients with high expression of each of the analyzed proteins (Extended Data Fig. 5). This analysis suggests that each of these proteins individually cannot predict suitability for eventual immunotherapies using γδ T cells.

Next, we aimed to elucidate the mechanisms involved in the killing of BCSC5 by γδ T cells, to narrow down which phenotypic changes might be mainly responsible for the immune escape of xenografted BCSC5. We investigated the role of Fas, MICB, TRAILR1/2 and ICAM-1 in the recognition and subsequent killing of BCSC5 by performing cytotoxicity experiments using blocking antibodies. We found that γδ T cell-mediated killing of BCSC5 involves Fas-FasL interactions, MICA/B engagement and, most drastically, ICAM-1 binding (Fig. 6d). In contrast, TRAIL-TRAILR interactions, ULBP2/5/6 engagement, and the binding of CD103 (integrin α E) did not play a major role in BCSC5 killing (Fig. 6d). These results indicate that among the phenotypic changes observed upon xenotransplantation, the loss of Fas, MICA/B and ICAM-1 expression might play a crucial role in protecting these cells from γδ T cell-mediated cytotoxicity. Our data also suggest that γδ T cells might require multiple receptor-ligand interactions to efficiently kill BCSC. We hence clustered TNBC patients based on the level of expression of the three key proteins Fas, MICB and ICAM-1. Cox proportional hazards analysis showed that patients expressing high levels of these proteins have a 1.4 times lower 5-year mortality risk (Fig. 6e).

### Zoledronate sensitizes Xenograft-derived tumor cells to γδ T cell-mediated killing

In addition to proteins directly involved in killing, we identified HMG-CoA reductase (HMGCR), the rate-limiting enzyme of the mevalonate pathway (Jo and DeBose-Boyd 2010; Friesen and Rodwell 2004) to be significantly downregulated in the xenograft-derived tumor cells (Fig. 6b and Fig. 7a). The mevalonate pathway regulates the levels of pAgs in tumor cells. These pAgs can be recognized by the Vγ9Vδ2 TCR expressed by Vδ2^+^ T cells in the context of butyrophilin 3A1 (BTN3A1) and 2A1 (BTN2A1) (Rigau et al. 2020; Karunakaran et al. 2020). Expression of BTN3A1, however, was not significantly altered in xenograft-derived tumor cells, and BTN2A1 was not detected in the proteomic study (Fig. 6b). Likewise, no changes were observed using an antibody recognizing all BTN3A isoforms (Fig. 6c). Considering that the BTN3A1 levels were comparable between BCSC5 culture cells and xenograft-derived tumor cells, we hypothesized that the loss of HMGCR expression was responsible for unresponsiveness of γδ T cells due to low pAg levels in the tumor cells. To further explore this recognition axis, we performed cytotoxicity assays in the presence of zoledronate or mevastatin, two well-studied drugs interfering with the mevalonate pathway and modulating pAg levels (Thompson et al. 2010; Zysk et al. 2018; Thompson and Rogers 2004; Li et al. 2009). The accumulation of pAgs by zoledronate increased γδ T cell cytotoxicity towards xenograft-derived tumor cells (Fig. 7b). The cytotoxicity in the presence of zoledronate was similar to the killing of untreated BCSC5 culture cells (Fig. 7b). In contrast, the accumulation of pAgs by zoledronate in BCSC5 culture cells failed to increase γδ T cell cytotoxicity towards these cells, suggesting that this recognition axis was already saturated (Fig. 7b). Reducing pAg levels by mevastatin led to reduced killing of BCSC5 culture cells and almost completely abolished the killing of xenograft-derived cells (Fig. 7b). These findings show that the killing of xenograft-derived tumor cells by γδ T cells almost exclusively relies on the recognition of pAgs and that these cells can be sensitized to γδ T cell-mediated killing by zoledronate treatment. Both BTN2A1 and BTN3A1 are necessary for the recognition of pAgs by γδ T cells (Karunakaran et al. 2020; Rigau et al. 2020). The correlation between the mRNA levels of BTN2A1 and BTN3A1 does not change between mammary healthy tissue and TNBC patient-derived samples (data not shown). We clustered TNBC patients by the expression levels of both BTN2A1 and BTN3A1 and found that high expression of both proteins significantly correlates with increased survival. Patients with high levels of BTN2A1 and BTN3A1 mRNA have a 2 times lower 10-year mortality risk (Fig. 7d). BCSC5 culture cells, in contrast, can still be partially recognized in the presence of mevastatin, in line with the finding that BCSC5 can be killed by several mechanisms, namely those involving Fas, MICB and ICAM-1 as detailed above (Fig. 7b and 6d). Thus, we clustered TNBC patients by the mRNA levels of BTN2A1 and BTN3A1, which are key for the recognition of differentiated cells, and Fas, MICB and ICAM-1, which seem to be central for the recognition and killing of BCSCs. Patients with high levels of these proteins showed significant increased survival having a 1.4 times lower 5-year mortality risk (Fig. 7e). Gene set enrichment analysis revealed that TNBC patients clustered by high (upper-quartile) average expression of BTN2A1, BTN3A1, Fas, MICB and ICAM-1 exhibited higher expression of BCSC genes and gene signatures associated to immune response, inflammation, IFN-γ and IFN-α responses, cytokine and chemokine signaling and immune-mediated cytotoxicity (data not shown) suggesting a favorable tumor immune microenvironment. To deepen into this observation, we applied the “deep deconvolution” CIBERSORT algorithm to deduce the immune cell composition (Newman et al. 2015) and compared the TNBC patients with high *versus* low average expression of BTN2A1, BTN3A1, Fas, MICB and ICAM-1 (Extended Data Fig. 6). CD4 memory T cells, resting and activated, CD8 T cells and anti-tumor M1 macrophages were enriched in TNBC patients with high average expression. In contrast, M2 tumor promoting macrophages were significantly reduced. The assessment of tumor infiltrating human γδ T cells by deconvolution has proven to be challenging, therefore we analyzed genes specifically upregulated in human γδ T cells that has been identified by machine learning approaches (Tosolini et al. 2017). Indeed, these human γδ T cell specific genes were clearly upregulated in those TNBC patients clustered by high (upper-quartile) average expression of BTN2A1, BTN3A1, Fas, MICB and ICAM-1 (Extended Data Fig. 6). Thus, expression of these proteins might be an instrumental tool to identify TNBC patients suitable for γδ T cell-based immunotherapy approaches.

**Fig. 7:**
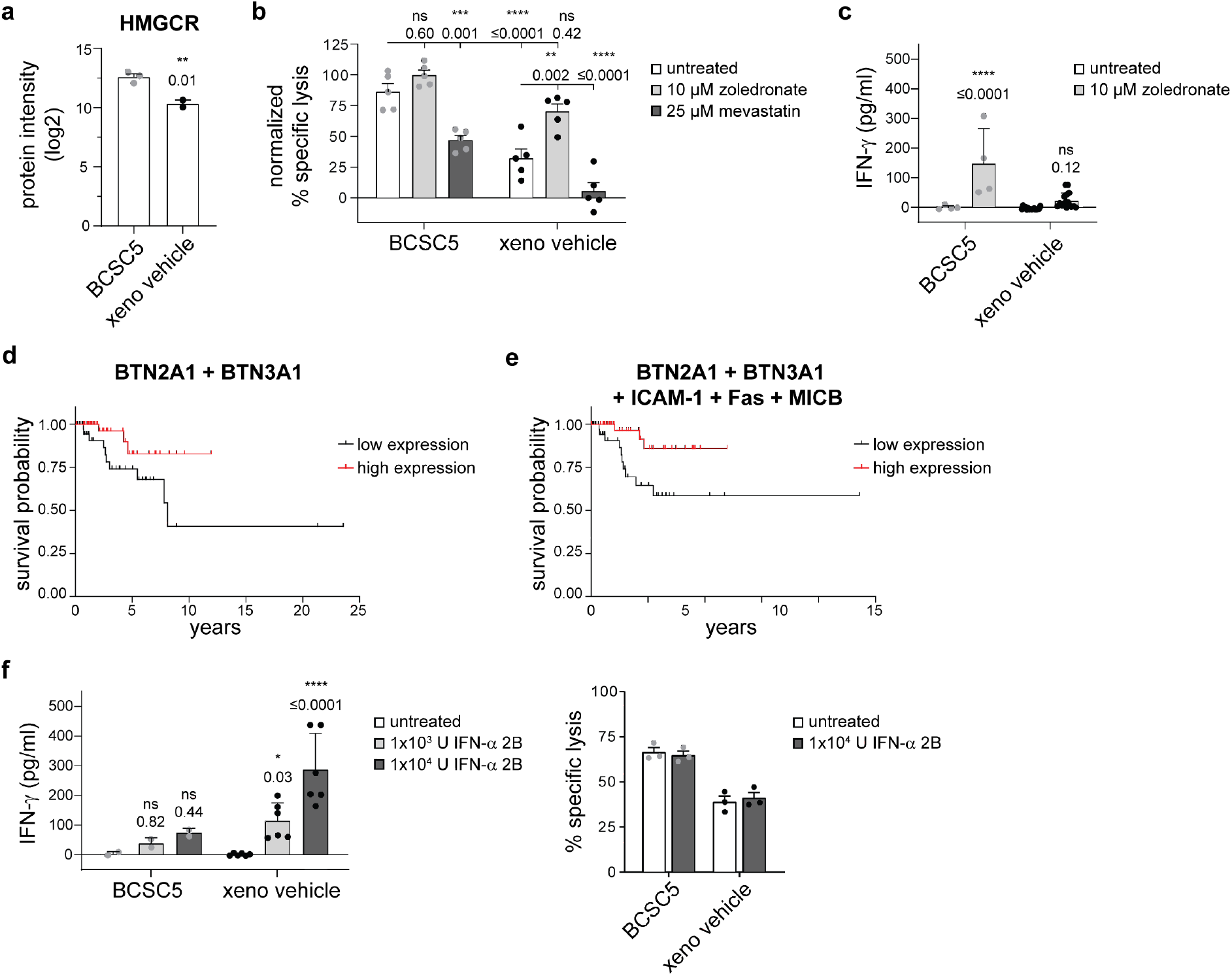
Zoledronate sensitizes Xenograft-derived tumor cells to γδ T cell-mediated killing. **(a)** HMGCR expression levels determined by mass spectrometry. Protein intensities (log2) are shown as means ± SEM for culture and xenograft-derived tumor cells (vehicle). Unpaired t-test, two-tailed. **(b)** *In vitro* killing of ^51^Cr-labeled tumor cells by γδ T cells. Target cells were pre-treated with 10 µM zoledronate or 25 µM mevastatin for 2 h. Cells were co-cultured at an E:T ratio of 10:1 for 20 h in the presence of the indicated drug. Results for three healthy donors of γδ T cells from four independent experiments were pooled (means ± SEM). Killing of BCSC5 culture cells in the presence of zoledronate was set to 100% for normalization. Two-way ANOVA followed by Tukey’s post hoc test comparing all groups to each other. **(c)** IFN-γ secretion by γδ T cells in response to BCSC5 culture cells or xenograft-derived tumor cells was determined. Target cells were pre-treated with 20 µM zoledronate for 2 h before co-culture with γδ T cells at a ratio of 1:1 in the presence of 10 µM zoledronate. The culture supernatant after 24 h was analyzed for secreted IFN-γ using ELISA. Results were baseline-corrected by the corresponding amounts of IFN-γ secreted by γδ T cells alone. Results for three healthy donors of γδ T cells from two experiments testing eight individual tumors were pooled (means ± SEM). Two-way ANOVA followed by Sidak’s post hoc test comparing treatment to respective untreated groups. **(d)** Cox regression of progression-free survival for TNBC patients sorted by high (upper-quartile) and low (lower-quartile) average expression of both proteins BTN2A1 and BTN3A1. β cox coefficient was -0.8 and the cox p value was 0.008. **(e)** Cox regression of progression-free survival for patients sorted by high (upper-quartile) and low (lower-quartile) average clustered expression of BTN2A1, BTN3A1, Fas, MICB and ICAM-1. β cox coefficient was -0.6 and the cox p value was 0.02. **(f)** IFN-γ secretion by γδ T cells in response to BCSC5 culture cells or xenograft-derived tumor cells was determined. Tumor cells were pre-treated with 10^3^ U or 10^4^ U IFN-α2B for 1 h before co-culture with γδ T cells at a ratio of 1:1 in the presence of IFN-α2B. The culture supernatant after 24 h was analyzed for secreted IFN-γ using ELISA. Results were baseline-corrected by the corresponding amounts of IFN-γ secreted by γδ T cells alone. Results for two healthy donors of γδ T cells from one experiment testing three individual tumors were pooled (means ± SEM). Two-way ANOVA followed by Dunnett’spost hoc test comparing treatment to respective untreated groups. **(e)** *In vitro* killing of ^51^Cr-labeled tumor cells by γδ T cells. Target cells were pre-treated with 10^4^ U IFN-α2B for 1 h and then co-cultured with γδ T cells in the presence of IFN-α2B at an E:T ratio of 10:1 for 20 h. Representative results for one healthy donor of γδ T cells is shown. Indicated are technical replicates (means ± SEM). * p≤0.05, ** p≤0.01, *** p≤0.001, **** p≤0.0001.

In addition to cytotoxicity, γδ T cells play a critical role in protective immune responses against tumor development by providing an early source of the pro-inflammatory cytokine IFN-γ (Chen et al. 2017; Gao et al. 2003). IFN-γ plays a manifold role in activating anticancer immunity. For instance, IFN-γ promotes the activity of tumor-triggered αβ T cells and inhibits the differentiation and activation of regulatory αβ T cells (Castro et al. 2018). The current view is that IFN-γ producing cells are endowed with potent cytotoxic functions during antitumor responses (Lopes and Silva-Santos 2021). In our settings, γδ T cells, with or without zoledronate treatment, did not respond with IFN-γ secretion to xenograft-derived tumor cells (Fig. 7c), which might limit the protective immune response against tumor development. It has been previously shown that IFN-α can mediate an increase in IFN-γ secretion by pAg-activated Vδ2^+^ T cells (Cimini et al. 2012). In line with this report, the pre-treatment of tumor cells with IFN-α induced γδ T cell-mediated IFN-γ secretion in response to xenograft-derived tumor cells (Fig. 7f) while failed to rescue γδ T cell-mediated cytotoxicity of xenograft-derived tumor cells (Fig. 7e). Taken together, the treatment with IFN-α facilitated the activation of γδ T cells to produce IFN-γ, which might be therapeutically interesting in clinical settings in which tumor-triggered αβ T cells play a significant role. Yet, this boost of γδ T cells to produce IFN-γ did not translate into increased cytotoxicity towards BCSCs highlighting that distinct signals and/or a different threshold of activation are needed to produce IFN-γ or to induce killing by γδ T cells. The molecular identification and targeting of the pathways regulating these two effector functions are beyond the scope of the present study.

In summary, our proteomic studies and biological validations strongly support the hypothesis that BCSC5 culture cells differentiated *in vivo*. This differentiation involved the upregulation of inhibitory T cell receptors, the loss of stem cell characteristics, the reduction of numerous γδ T cell ligands and a decrease in pAg levels, in sum preventing efficient recognition and killing by γδ T cells.

## Conclusions

It has been suggested that BCSCs are responsible for therapy resistance and metastatic dissemination in breast cancer, which is the leading cause of cancer deaths among women worldwide (Sung et al. 2021). Until now, treatment resistant BCSCs have only been poorly characterized, and targeted therapeutics have yet to be identified. We have recently established an optimized culture system to expand human BCSCs that faithfully reproduce the original patient’s tumor characteristics and are therefore an ideal cellular platform to test novel therapeutics (Metzger et al. 2017; Strietz et al. 2021). Here, we aimed to investigate the susceptibility of these BCSCs to immunotherapy using primary human γδ T cells. We showed that patient-derived triple negative BCSCs are targeted by both Vδ1^+^ and Vδ2^+^ primary expanded γδ T cells. However, orthotopically xenografted BCSC5, the BCSC line best recognized by γδ T cells in our study, was refractory to γδ T cell immunotherapy. We demonstrated that attraction and/or migration towards xenografted cells was not the reason for this unexpected *in vivo* outcome. Both Vδ1^+^ and Vδ2^+^ T cells migrated efficiently towards BCSC-conditioned medium and this migration was even increased towards xenografted cells. The chemokine receptors expressed on the expanded γδ T cells were compatible with the secretion profile of xenografted cells.

Solid tumors, including breast cancer, are often surrounded by a dense ECM preventing the efficient infiltration of immune cells (Goetz et al. 2011; Cohen and Blasberg 2017). A high stromal content has been associated with poor prognosis in TNBC, and an accumulation of γδ T cells in the tumor stroma contributes to this observation (Hidalgo et al. 2014; Kramer et al. 2019). Detailed analyses of γδ T cell migration with respect to the tumor stroma are not yet available. Here, we found that exogenously expressing MMP14 conferred pro-migratory functions to primary γδ T cells in a 3D model for basement membranes. Crossing of the basement membrane is of critical relevance for γδ T cells to extravasate from the blood stream at the tumor site (Slaney et al. 2014; Donnadieu et al. 2020). Using viable slices of BCSC5-derived xenograft tumors, we showed that γδ T cell migration was accelerated inside of tumor islets compared to the ECM-rich tumor stroma similar to previous observations made for αβ T cells (Bougherara et al. 2015). MMP14 expression increased γδ T cell migration speed exclusively in the tumor stroma and not inside the tumor tissue, supporting the functional role of MMP14 in cleaving the peri-tumoral ECM (Pei and Weiss 1996; Ohuchi et al. 1997). These findings are in line with a previous report showing that expressing the secreted ECM-modifying enzyme heparanase in human CAR αβ T cells promoted tumor rejection in melanoma and neuroblastoma xenograft models (Caruana et al. 2015). In another study, inhibition of the ECM-crosslinking enzyme lysyl oxidase improved anti-tumor responses in combination with checkpoint inhibition due to increased αβ T cell migration and infiltration (Nicolas-Boluda et al. 2021). We hypothesize that the overexpression of MMP14 might have certain advantages over the expression of secreted heparanase or the systemic application of lysyl oxidase inhibitors, as MMP14 is a membrane-anchored protein lowering the risk of structural changes outside of the tumor tissue. Although we have not observed any side effects related to MMP14 expression in mice receiving γδ T cell treatment, the application of MMP14 will have to be strictly monitored with respect to biodistribution and its impact on healthy tissues in future studies.

Despite increased chemoattraction and MMP14-mediated peri-tumoral migration, γδ T cells failed to control BCSC5-derived tumors *in vivo*. Remarkably, xenografted cells showed reduced capacity to activate γδ T cells and thereby reduced susceptibility to γδ T cell-mediated cytotoxicity. Despite our BCSCs faithfully reproducing many original patient’s tumor characteristics (Metzger et al. 2017; Strietz et al. 2021), they underwent major changes in their proteomic signature after xenograft and growth *in vivo*. BCSC5 cells lost their stem cell characteristics and downregulated a plethora of surface proteins key for immunosurveillance by γδ T cells after transplantation and growth *in vivo.* Intriguingly, this *in vivo* differentiation was not a consequence of mechanisms induced by the immune system or the immunotherapy, since it was equally observed in immunodeficient mice with or without the presence of primary human γδ T cells. Xenografted cells exhibited increased surface expression of the inhibitory T cell ligands PD-L1 and PD-L2 compared to parental BCSC5 cells, in line with studies associating TNBC with high expression levels of the PD-L1 (Mittendorf et al. 2014; Nanda et al. 2016; Li et al. 2021). This increased expression of inhibitory T cell ligands might explain the reduced susceptibility to be killed by γδ T cells. However, a combinatorial treatment of xenografts with γδ T cells and α-PD-1 did not result in reduced xenograft growth. Similarly, Li and colleagues have described the inefficacy of this treatment regimen against TNBC MDA-MB-231-derived xenografts (Li et al. 2021).

Because increased PD-L1 levels were not responsible for the immune evasion of xenografted cells, we searched for additional molecules involved in BCSC recognition and compared their expression levels before and after *in vivo* growth. Our results revealed that recognition and killing of BCSC5 by γδ T cells required the engagement of multiple receptor-ligand pairs. Killing was thus dependent on ICAM-1 binding, MICB, pAgs/BTN2A1/BTN3A1 and on Fas/FasL interactions. Indeed, clustering TNBC patients for high expression of these proteins significantly increased survival prognosis. In αβ T cells, ICAM-1 is key in the formation of a functional immune synapse (Grakoui 1999) and to enable CAR αβ T-cell entry into solid tumors (Kantari-Mimoun et al. 2021). Low ICAM-1 expression levels on breast cancer cells made them resistant to αβ T cell killing (Chen et al. 2017). Likewise, human pancreatic cancer cell lines lacking ICAM-1 were poorly bound and killed by γδ T cells *in vitro* (Liu et al. 2009). Thus, our results support that ICAM-1-mediated intercellular interactions facilitate γδ T cell-mediated recognition and killing of BCSCs by Fas/FasL interactions and pAgs.

However, xenograft-derived tumor cells lost Fas, MICB and ICAM-1 expression, and downmodulated HMGCR, the rate-limiting enzyme of the mevalonate pathway producing pAgs. These changes are most probably responsible for the escape of xenograft-derived tumor cells from γδ T cell recognition. Yet we observed some residual cytotoxicity towards xenograft-derived tumor cells that was exclusively dependent on pAg recognition as pharmacological inhibition of HMGCR by mevastatin abolished cytotoxicity. Accumulation of pAgs by zoledronate pre-treatment overcame the resistance of xenograft-derived tumor cells to γδ T cell-mediated killing. The clinical application of zoledronate is FDA-approved for the treatment of osteoporosis and bone metastasis. Therefore, combinatorial therapy approaches using zoledronate and the adoptive transfer of γδ T cells represent a promising option to simultaneously tackle BCSCs and their differentiated progeny.

Our observation that BCSCs can be better recognized by γδ T cells than their differentiated progeny apparently opposes previous reports using the expression of CD44 and CD24 to define stem-like cells (CD44^hi^CD24^lo^) and their non-stem cell counterparts (CD44^hi^CD24^hi^). Sorted stem-like cells from the triple negative SUM149 cell line and from PDX401 cells were less efficiently killed by γδ T cells compared to non-stem cells counterparts (Dutta et al. 2021). This study identified MICA shedding from the tumor cell surface as a mechanism to escape γδ T cell recognition. MICA surface levels were unchanged between the differentiated xenograft cells and the BCSCs suggesting that MICA shedding does not play a major role in the escape of BCSC5-derived xenografts *in vivo*. To the best of our knowledge, only one other study investigated the response of γδ T cells against BCSCs and non-stem cells derived from *ras*-transformed human mammary epithelial cells, and found that both cell populations were equally resistant to γδ T cell-mediated killing (Chen et al. 2017). Yet, both populations could be sensitized by zoledronate treatment in line with our results. Taken together, these discrepancies among studies underline the importance of perceiving immunotherapeutic approaches as individualized medicine, which might have to be tailored for each patient.

In addition, our results highlight a previously unnoticed dichotomy, namely that IFN-γ responses and cytotoxicity by γδ T cells do not necessarily correlate. Zoledronate enhanced cytotoxicity by γδ T cells but failed to promote IFN-γ secretion. In contrast, IFN-α treatment increased γδ T cell-mediated IFN-γ secretion but failed to enhance γδ T cell-mediated cytotoxicity. This result suggests that IFN-α might represent an attractive tool to be used in combinatorial therapies to induce synergistic effects of γδ and αβ T cell responses.

In summary, our data show that patient-derived triple negative BCSCs are targetable by expanded γδ T cells. However, *in vivo* growth of these BCSCs leads to their differentiation into cells that lost stemness and ligands to activate γδ T cell responses and thereby, escaped from efficient killing by γδ T cells. Still, γδ T cells residually killed *in vivo* differentiated cells by recognizing pAgs. This killing could be increased to the level of BCSC5 killing by zoledronate. In all, we propose that a combinatorial therapy using γδ T cells and zoledronate represents a valuable approach to target triple negative BCSCs and non-stem cells alike. Furthermore, IFN-α treatment could induce IFN-γ production by γδ T cells, and thereby induce a first source of IFN-γ promoting ICAM-1 expression, T cell entry into solid tumors (Kantari-Mimoun et al. 2021) and further endogenous αβ T cell responses in immunocompetent settings.

## Methods

### Cell lines

Healthy human dermal fibroblasts of neonatal origin (HDFn) were cultured in DMEM GlutaMAX medium (Thermo Fisher) supplemented with 10% fetal bovine serum (FBS), 100 µg/ml penicillin, 100 µg/ml streptomycin and 10 mM HEPES (all Thermo Fisher). HEK293T cells were grown in DMEM GlutaMAX medium supplemented with 10% FBS, 100 µg/ml penicillin, 100 µg/ml streptomycin, 10 mM HEPES and 10 µM sodium pyruvate (Thermo Fisher). HDFn and HEK293T cells were cultured at 37°C in a humidified atmosphere of 7.5% CO_2_.

BCSCs were cultured in mammary stem cell (MSC) medium: MEBM medium (Lonza) supplemented with 1x B27 (Thermo Fisher), 1x amphotericin (Sigma Aldrich), 20 ng/ml EGF (Peprotech), 20 ng/ml FGF (Peprotech), 4 µg/ml heparin (Sigma Aldrich), 35 µg/ml gentamicin (Thermo Fisher), 500 nM H-1152 (Calbiochem), 100 µg/ml penicillin and 100 µg/ml streptomycin. Passaging of BCSCs was performed as previously described (Metzger et al. 2017). BCSCs were cultured at 37°C under low oxygen conditions (3% O_2_, 5% CO_2_, 92% N_2_).

### γδ T cell expansion

Primary γδ T cell expansion was performed as previously described (Siegers et al. 2013). Briefly, peripheral blood mononuclear cells (PBMCs) were purified from the blood of healthy donors via density gradient centrifugation (Pancoll human, Pan Biotech). PBMCs were resuspended at a concentration of 1x10^6^ cells/ml in γδ T cell medium (RPMI 1640 medium (Gibco) supplemented with 10% FBS, 100 µg/ml penicillin, 100 µg/ml streptomycin, 10 mM HEPES, 10 µM sodium pyruvate and 1x MEM non-essential amino acids (Pan Biotech)). Expansion was induced with 1 µg/ml ConcanavalinA (Sigma Aldrich), 10 ng/ml IL-2 (Peprotech) and 10 ng/ml IL-4 (Peprotech). Cells were adjusted every 3-4 days to 1x10^6^ cells/ml with γδ T cell medium and cytokines. The cells were cultured at 37°C in a humidified atmosphere of 5% CO_2_. At day 10-14 after expansion start, γδ T cells were separated from αβ T cells via negative selection (MACS TCRγ/δ^+^ T Cell Isolation Kit, Miltenyi Biotech). Cultures with a purity of ≥95% were used for experiments from day 14 on. Buffy coats used as blood source were purchased from the blood bank of the University Medical Centre Freiburg (approval of the University Freiburg Ethics Committee: 147/15).

### Lentiviral constructs, generation of lentiviral particles and transduction of γδ T cells

*MMP14* cDNA or the cDNA of the catalytically inactive form of MMP14 *(MMP14^E240A^*) (both kind gifts from Pilar Gonzalo and Joaquin Teixidó (Bartolomé et al. 2009)) were cloned into the lentiviral backbone pLVX-CMV-IRES-zsGreen1 (Takara/Clontech #632187) by Gibson assembly (Gibson et al. 2009). The CMV promoter was exchanged with a short EF-1α (sEF-1α) promoter sequence. The integrity of each plasmid was verified by restriction enzyme digestion and Sanger sequencing.

For the generation of lentiviral particles, 1x10^7^ HEK 293T cells were plated on 15 cm dishes and cultured at 37°C and 7.5% CO_2_. After 24 h, the medium was exchanged and HEK293T cells were transfected with the indicated constructs and the packaging plasmids pCMV-dR8.74 and pMD2-vsvG (both kind gifts from Didier Trono) using PEI transfection. After 24 and 48 h, the viral particle-containing supernatant was harvested, pooled, filtered and concentrated using density centrifugation (10% sucrose w/v in PBS/0.5  mM EDTA) for 4 h at 10,000xg and 6°C. The supernatant was discarded, and the viral particles were resuspended in PBS using 1/400^th^ of the harvested volume and stored at -80°C.

γδ T cells were lentivirally transduced with a multiplicity of infection (MOI) of 2-5 as indicated for the individual experiments. Transduced γδ T cells were checked for zsGreen1 and MMP14 expression by flow cytometry after 2-3 days post transduction.

### Flow cytometry analysis

To stain cell surface proteins, cells were washed once with flow cytometry buffer (PBS supplemented with 2% FBS) and incubated in the diluted antibody solution for 15 min at 4°C. In the case of fluorophore-labeled antibodies, cells were washed once with flow cytometry buffer and analyzed on a Gallios flow cytometer (Beckman Coulter). Unlabeled primary antibodies were visualized using fluorescently labeled secondary antibodies. After washing away the primary antibody, cells were incubated in the diluted secondary antibody solution for 15 min at 4°C. Finally, cells were washed once as described above and analyzed on a Gallios flow cytometer.

### Antibodies and chemicals

Self-made α-CD3ε (clone UCHT1) and α-CD28 (clone CD28.2) from BioLegend were used for γδ T cell stimulation.

For flow cytometry, the following antibodies were used: α-EpCAM-AlexaFluor488 (clone 9C4), α-CD107a-PE (clone H4A3), α-BTN3A-PE (clone BT3.1), α-Fas-BV421 (clone DX2), α-TRAILR1-APC (clone DJR1), α-TRAILR2-PE (clone DJR2-4), α-ICAM-1-BV421 (clone HA58), α-CD3ε-AlexaFluor488 (clone UCHT1), α-CD3ε-AlexaFluor647 (clone UCHT1), α-CD27-PE (clone M-T271), α-NKG2D-APC (clone 1D11), α-LAG-3-AlexaFluor647 (clone 11C3C65), α-PD-L1-APC (clone 29E.2A3), α-PD-L2-BV421 (clone 24F.10C12), α-Vδ2TCR-Biotin (clone B6), α-CXCR1-APC (clone 8F1), α-CXCR3-AlexaFluor647 (clone G025H7), α-CXCR4-PE (clone 49801), α-CXCR5-APC (clone J252D4), α-CXCR6-PE (clone K041E5), α-CCR2-APC/Fire 750 (clone K036C2), α-CCR3-FITC (clone 5E8), α-CCR10-APC (clone 6588-5) and α-TIM-3-PE-Cy7 (clone F38-2E2) from BioLegend. α-ULPB2/5/6 (mouse, clone 165903) and α-CCR4-APC (clone 205410) from R&D Systems. α-mouse IgG-APC from Southern Biotech. a-γδTCR-PE (clone SA6.E9), a-γδTCR-FITC (clone SA6.E9), α-rabbit-DyLight633 (polyclonal), α-PD-1-PE-Cy7 (clone J105), streptavidin-eFluor450 and streptavidin-PE-Cy7 from Thermo Fisher. α-CCR5-PE (clone 2D7), α-CCR6-BB515 (clone 11A9), α-CCR7-AlexaFluor647 (clone 150503) and α-CD45RA-V450 (clone HI100) from BD Bioscience. α-Vδ1TCR-APC (clone REA173) and α-Vδ1TCR-PE (clone REA173) from Miltenyi. α-MMP14 (rabbit, polyclonal) from Abcam. The Cell Proliferation Dye eFluor450 was purchased from Thermo Fisher.

α-TCRVδ1-Biotin (clone REA173) from Miltenyi and α-TCRVδ2-Biotin (clone B6) from BioLegend were combined with the antibody solution provided in the MACS TCRγ/δ^+^ T Cell Isolation Kit to separate Vδ2^+^ and Vδ1^+^ cells from γδ T cell expansion cultures, respectively. The following antibodies were used for immunofluorescence: α-EpCAM-BV421 (clone EBA-1) from BD Biosciences, α-fibronectin (rabbit, polyclonal) and α-rabbit IgG-AlexaFluor546 (polyclonal) from Sigma-Aldrich. CellTracker Green CMFDA Dye was used from Thermo Fisher.

The following antibodies were used for blocking experiments: α-Fas (clone A16086F), α-FasL (clone NOK-1), α-TRAIL (clone RIK-2), α-TRAILR1 (clone DJR1), α-TRAILR2 (clone DJR2-4), α-NKG2D (clone 1D11), α-MICA/B (clone 6D4), α-ICAM-1 (clone HCD54), α-IgG1 isotype (clone MOPC-21) and α-IgG2b isotype (clone MG2b-57) from BioLegend. α-ULPB2/5/6 (clone 165903) from R&D Systems. α-CD103 (clone 2G5) from Beckman Coulter. Zoledronic acid monohydrate (zoledronate) and mevastatin were purchased from Sigma-Aldrich.

Viability of γδ T cells was assessed using the FITC Annexin V Apoptosis Detection Kit I (BD Bioscience).

### Cytotoxicity and degranulation assay

Bioluminescence-based cytotoxicity assays were performed as previously described (Karimi et al. 2014). Briefly, 1x10^4^ luciferase-expressing target cells were plated in a white 96-well flat bottom plate. Effector cells were added to the target cells at the desired effector to target cell ratios. 37.5 µg/ml D-Luciferin Firefly (Biosynth) was added to the samples, which were then incubated for the indicated time at 37°C and measured using a luminometer (Tecan infinity M200 Pro). Bioluminescence was measured as relative light units (RLUs). RLU signals from target cells alone served as spontaneous death controls. Maximum killing RLUs were determined using target cells lysed with 1% Triton X-100. The specific lysis was calculated with the following formula:

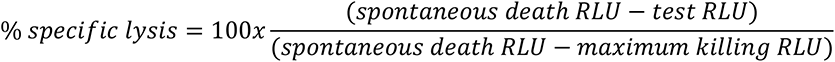

^51^Cr-release assays were performed to assess the cytotoxicity in experiments including xenograft-derived tumor cells. Target cells (tumor cells) were loaded with ^51^Cr for 1 h at 37°C. After washing the cells 3 times with medium, cells were resuspended in γδ T cell medium and 1x10^4^ cells were plated in a 96-well U bottom plate. Effector cells (γδ T cells) were added to the target cells at the desired effector to target cell ratios. Samples were incubated for the indicated time at 37°C. Supernatants were transferred to a solid scintillator-coated 96-well plate (LumaPlate, Perkin Elmer) and then measured using a microplate scintillation γ-ray counter (TopCount, Perkin Elmer). Target cells without effector cells served as spontaneous ^51^Cr release controls. Maximum ^51^Cr release was determined using target cells lysed with 1:20 centrimide. The specific lysis was calculated with the following formula:

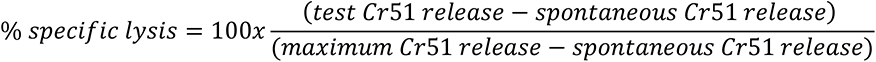

For blocking experiments, effector or targets cells were pre-incubated with 20 µg/ml of the indicated blocking antibodies at 37°C for 1 h. IgG isotype antibodies were used as experimental controls. Cells were then used for cytotoxicity experiments in the presence of 10 µg/ml of the blocking antibodies as detailed above.

The blocking agent mevastatin was pre-incubated with the target cells at a concentration of 25 µM for 2 h at 37°C. The cytotoxicity assay was conducted in the presence of 25 µM mevastatin. Experiments with zoledronate were performed depending on the target cells. BCSC culture cells were pre-incubated with 10 µM zoledronate over night at 37°C and washed before the cytotoxicity assay. In contrast, for experiments including xenograft-derived tumor cells, all target cells were pre-incubated with 10 µM zoledronate for 2 h and the cytotoxicity assay was performed in the presence of 10 µM zoledronate.

To analyze the effect of IFN-α 2B, target cells were pre-incubated with 2x10^3^ U or 2x10^4^ U IFN-α 2B (Stemcell) for 1 h at 37°C. The assays were conducted with a final concentration of 1x10^3^ U or 1x10^4^ U IFN-α 2B.

To analyze γδ T cell degranulation, BCSCs or xenograft-derived tumor cells were labeled with 20 µM cell proliferation dye eFluor450 (Thermo Fisher). 1x10^5^ tumor cells and 1x10^5^ γδ T cells were co-cultured for 3 h at 37°C in the presence of 1 µl α-CD107a-PE antibody (BD Biosciences). Medium or stimulation with α-CD3 and α-CD28 (both 1 µg/ml; plate-bound) served as negative and positive controls, respectively. Cells were harvested and analyzed by flow cytometry.

### IFN-γ ELISA

1x10^5^ γδ T cells were co-cultured with tumor cells for Target cells were pre-incubated with 20 µM zoledronate for 2 h or with 2x10^3^ U or 2x10^4^ U IFN-α 2B for 1 h at 37°C before co-culturing them with γδ T cells. The assay was conducted with a final concentration of 10 µM zoledronate or 1x10^3^ U or 1x10^4^ U IFN-α 2B. The culture supernatant was then analyzed for secreted IFN-γ was detected using an IFN-γ ELISA kit (Thermo Fisher) according to the manufacturer’s instructions.

### Xenograft tumor model and γδ T cell treatment

NOD SCID mice (NOD.CB17-Prkdcscid/Rj) were purchased from Janvier Labs and housed at the Center for Experimental Models and Transgenic Service, Freiburg, under specific pathogen-free conditions using individually ventilated cages. Mouse handling and experiments were performed in accordance with German Animal Welfare regulations and approved by the local authorities (animal protocols G17/137).

The orthotopic transplant was performed as described previously (Metzger et al. 2017). 5x10^5^ BCSCs were mixed with 1x10^6^ irradiated HDFn in a 1:1 mixture of MSC medium and Matrigel (growth factor reduced, Corning) and transplanted into each fat pad of the two #4 mammary glands of a female NOD SCID mouse. Mice were anesthetized during the procedure using an isoflurane inhalator. Before treatment start, mice were randomized into three groups: vehicle (PBS), γδ T cell, γδ T cell MMP14. Treatment was initiated for each mouse individually at a tumor diameter of at least 4 mm^3^ as indicated for each experiment. 5x10^6^ γδ T cells (culture purity ≥95%; <5% Vδ1^+^) were intravenously injected three times per week for the indicated time period. In addition, mice received 0.6x10^6^ IU of IL-2 (Proleukin, Novartis) in incomplete Freund’s adjuvant subcutaneously in the abdomen to support γδ T cell survival *in vivo*. Checkpoint inhibition using the α-PD-1 antibody Nivolumab (Opdivo, Bristol Myers Squibb) and a corresponding α-human IgG4κ isotype control (Crown Bioscience) was performed by additional biweekly intraperitoneal injections of 200 µg of the respective antibody. Tumor sizes were defined by caliper measurement. Tumor volumes were calculated using the formula V = 4/3 x π x r^3^.

### Preparation of tumor single cell suspensions

Tumors from BCSC5 orthotopic xenografts were cut into small pieces using a razor blade and digested with 1 mg/ml collagenase IV (Sigma Aldrich) and 0.1 mg/ml DNAse I (Roche) at 37°C for 45 min. The digested tissue was filtered through 70 and 40 µm filters. To remove remaining cell debris, the cell suspension was cleared via centrifugation through a FBS layer. Erythrocytes were removed using Ammonium-Chloride-Potassium (ACK) lysis and recovered cells were used for experiments. For flow cytometric analysis, EpCAM-positive cells were gated.

### Preparation of murine blood and liver samples

Leukocytes were isolated from blood samples by repeated erythrocyte lysis steps using ACK lysis buffer. When only minimal residual erythrocytes were left in the sample, cells were used for flow cytometric analysis. To analyze liver-derived lymphocytes, liver tissue was cut into small pieces using scissors and further dissociated through a 70 µm filter. The cell suspension was centrifuged at 60xg for 1 min at room temperature (RT) without break and the resulting supernatant was centrifuged at 850xg for 8 min at RT. The pellet was then resuspended in 10 ml 37.5% Percoll (Sigma Aldrich) in PBS and 100 U/ml heparin (Sigma Aldrich) and centrifuged at 850xg for 20 min at RT without break. Erythrocytes were removed from the pellet using ACK lysis and recovered cells were used for flow cytometry.

### Preparation of viable BCSC5 xenograft tumor slices and confocal imaging of γδ T cell migration

Tumor slices were prepared as described previously (Salmon et al. 2011; Salmon et al. 2012). Briefly, BCSC5 xenograft-derived tumors were cut into small pieces using a razor blade. Tumor pieces were embedded in a 5% low-gelling-temperature agarose (Sigma Aldrich) solution (w/v in PBS). After agarose solidification at 4°C, agarose blocks were fixed on the specimen disk of a vibratome using non-toxic tissue adhesive (3M Vetbond). The embedded tissue was cut into 350 µm thick slices in ice-cold PBS. Tumor slices were transferred onto 30 mm organotypic culture insert (Merck), which had been placed in the wells of a 6-well plate filled with 1.1 ml phenol red-free RPMI medium (Thermo Fisher) supplemented with 10% FBS, 100 µg/ml penicillin and 100 µg/ml streptomycin (slice assay medium).

For fluorescent labeling of the tumor tissue and plating of γδ T cells, pre-wet stainless steel flat washers were placed onto the agarose surrounding each tumor slice and the slices were incubated at 37°C for 10 min. Then, tumor slices were stained with α-EpCAM-BV421 (10 µg/ml) and α-fibronectin (3.5 µg/ml) antibodies for 15  min at 37°C and subsequently washed with slice assay medium. 1x10^6^ γδ T cells were labeled with 0.5 µM CellTracker Green CMFDA Dye (Thermo Fisher) according to the manufacturer’s instruction and mixed with an α-rabbit-AlexaFluor564 antibody (10 µg/ml). The solution was added on the tumor slices and incubated for 30 min at 37°C. Subsequently slices were washed and incubated at 37°C for 10 min until imaging.

Image acquisition was performed at 37°C in slice assay medium with a LSM 880 inverted laser scanning confocal microscope (Zeiss) equipped with a 25x objective (LD LCI Plan-Apochromat, NA 0.8, WD 0.57 mm, water immersion) using the Fast Airyscan mode. BV421, CMFDA and AlexaFluor546 were excited with a 405, 488 or 561 nm laser respectively. Nine optical planes spanning a total depth of 63 µm in the Z dimension were captured every 30 s for 20-45 min.

Airyscan data were first processed and stitched using the Zeiss Zen Black edition 3.0 SR. Then, ECM regions were manually defined with the help of the fibronectin staining in each individual plane along the Z axis. Areas negative for fibronectin signal were considered as tumor regions. Further data analysis was performed using Python and the scikit-image library (van der Walt et al. 2014). Images were first corrected for sample drift by detecting matching features in subsequent frames of the ECM channel and estimating transformation parameters based on their coordinates. Cells were segmented by intensity after background correction to reduce bleed-through of signal from the ECM channel and median filtering for smoothing. Detected features were filtered by size to remove noise and cell clumps. To facilitate tracking, cells that were apparent in more than one Z plane were only considered in the plane in which their size was maximal. Subsequent tracking of the detected cells was performed using Trackpy (Allan et al. 2019), and two tracks were merged if their respective initial or final point were less than 10 µm and less than two frames apart. Only tracks with data for at least 6 frames were considered for further analysis.

For localization analysis, cells were considered to localize to tumor regions when no pixel of the γδ T cell CMFDA signal overlapped with the fibronectin signal. A cell was defined to have a tumor dwell time >50% if it spent at least 50% of its observed frames localized to tumor regions. For speed analysis, the momentary speed of a cell was defined as the distance travelled between two successive observations of the same cell, divided by the time between these observations.

Representative microscopy images and videos were generated using Imaris 9.3.1.

### Preparation of tumor cell-derived conditioned medium

For transwell migration assays, conditioned medium (CM) was generated by harvesting the supernatant of BCSCs cultures after 5 days. Culture supernatants were cleared from residual cells by centrifugation and stored at -80°C until use.

To generate CM from xenograft-derived cells, 5x10^6^ tumor cells were cultured in γδ T cell medium for 24 h at 37°C in a humidified atmosphere of 5% CO_2_. Culture supernatants were cleared from residual cells by centrifugation and directly used for experiments.

### Transwell and Matrigel migration assays

For transwell migration assays towards CM, the CM was diluted 1:1.67 in γδ T cell medium/1% FBS. 250 µl were transferred to the receiver wells of a 96-well transwell plate (Corning). Medium without cells, incubated in the same conditions as BCSCs, served as negative control. For migration towards the chemokine CXCL12, γδ T cell medium/1% FBS was supplemented with 50 ng/ml CXCL12.

For transwell assays towards tumor cells, tumor cells were labeled with 20 µM cell proliferation dye eFluor450 (Thermo Fisher). 1.5x10^5^ tumor cells in 250 µl γδ T cell medium/1%FBS were seeded into the receiver wells of a 96-well transwell plate. γδ T cell medium/1%FBS served as negative control.

2x10^5^ γδ T cells in 100 µl γδ T cell medium/1% FBS were seeded in the well of the permeable support with 5 µm pore size (Corning). After 3 h incubation at 37°C, transmigrated cells were stained to distinguish T cell subsets and counted via flow cytometry. Matrigel migration assays were performed as previously described (Goetz et al. 2011; Hörner et al. 2019). Briefly, 3.5x10^4^ γδ T cells were resuspended in 10 µl ice-cold Matrigel (Corning) and plated into the inner well of a pre-cooled µ-Slide Angiogenesis (ibidi). After cell settling at 4°C, samples were solidified at 37°C. 15 µl Matrigel were added on top of the cell-containing gels. Following solidification at 37°C, wells were filled with γδ T cell medium supplemented with 10% FBS, 30 U/ml IL-2 and 50 ng/ml CXCL12 (Peprotech). After 48 h, samples were fixed with 4% PFA/0.25% glutaraldehyde for 20 min at RT. Gels were washed with PBS and cells were permeabilized with 0.5% Triton-X100 (Carl Roth) in PBS for 40 min. After washing, nuclei were stained with 5 µg/ml Hoechst 33342 (Sigma Aldrich) for 3 h in the dark and samples were imaged using confocal microscopy.

Images were acquired using a C2 confocal microscope (Nikon) with a 20x objective (NA: 0.75, WD: 1 mm). Hoechst 33342 was excited with a 405 nm laser. Z stacks with a step size of 2 µm were imaged. Quantification of the migration distance was performed automated by using Python and the scikit-image library (van der Walt et al. 2014). Briefly, the voxels of the stacks were segmented into foreground (cells) and background by intensity thresholding. Overlapping cells were then separated in 3D with the Watershed algorithm. The position of a cell was defined to coincide with its centroid. Migration distances were normalized between images by baseline subtraction per image: the baseline migration distance for an image was defined as the 10^th^ percentile of all cells’ Z coordinates. This value was subtracted from all Z coordinates. Z coordinates were set to 0 if they became negative after baseline correction.

### Proteomics

#### Sample preparation

Proteomic studies were performed following previously published protocols (Hughes et al. 2014). BCSC5 culture cells and xenograft-derived tumor cells were washed with PBS, pelleted, frozen in liquid nitrogen and stored at -80°C until further processing. Cell pellets were lysed in 400 µl lysis buffer (4% sodium dodecyl sulfate, 50 mM tetraethylammonium bromide (pH 8.5) and 10 mM tris(2-carboxyethyl)phosphine hydrochloride). Lysates were boiled for 5 min and then sonicated for 15 min at high intensity (30 sec on /30 sec off). After sonication, DNA and RNA were degraded using Benzonase endonuclease (Merck Millipore). The protein concentration was measured with EZQ Protein Quantitation Kit (Thermo Scientific). Lysates were alkylated in the dark with 20 mM iodoacetamide for 1h at RT. For protein clean-up, 200 µg SP3 paramagnetic beads were added to the lysates, and proteins were bound to the beads by adding acetonitrile with 0.1% formic acid. Beads were washed in 70% ethanol and 100% acetonitrile before elution in digest buffer (0.1% sodium dodecyl sulfate, 50 mM tetraethylammonium bromide (pH 8.5) and 1 mM CaCl_2_) and digested sequentially with LysC (Wako), then Trypsin (Promega), each at a 1:100 w/w (enzyme:protein) ratio. Peptide clean-up was performed according to the SP3 protocol.

### Tandem mass tag (TMT) labelling and basic C18 reverse phase (bRP) chromatography fractionation

Each sample (200 µg of peptides each) was resuspended in 100 µl of 100 mM tetraethylammonium bromide buffer. TMT-10plex (Thermo) labelling was performed according to manufacturer’s protocol. To ensure complete labelling, 1 µg of labelled samples from each channel was analysed by LC-MS/MS prior to pooling. The mixture of TMT 10plex sample was desalted with Sep Pak C18 cartridge (Waters), and then fractionated by basic C18 reverse phase chromatography as described (Roth et al 2020 10.1016/j.chembiol.2020.06.012)

### Liquid chromatography electrospray tandem mass spectrometry analysis (LC MS/MS analysis)

The LC separations were performed as described (Roth et al) with a Thermo Dionex Ultimate 3000 RSLC Nano liquid chromatography instrument. Approximately 1 µg of concentrated peptides (quantified by NanoDrop) from each fraction were separated over an EASY Spray column (C18, 2 µm, 75 µm × 50 cm) with an integrated nano electrospray emitter at a flow rate of 300 nL/min. Peptides were separated with a 180 min segmented gradient. Eluted peptides were analysed on an Orbitrap Fusion Lumos (Thermo Fisher) mass spectrometer.

### Data Analysis

All the acquired LC-MS data were analysed using Proteome Discoverer software v.2.2 (Thermo Fisher) with Mascot search engine. A maximum missed cleavages for trypsin digestion was set to 2. Precursor mass tolerance was set to 20 ppm. Fragment ion tolerance was set to 0.6 Da. Carbamidomethylation on cysteine and TMT-10plex tags on N termini as well as lysine (+229.163 Da) were set as static modifications. Variable modifications were set as oxidation on methionine (+15.995 Da) and phosphorylation on serine, threonine, and tyrosine (+79.966 Da). Data were searched against a complete UniProt Human (Reviewed 20,143 entries downloaded Nov 2018). Peptide spectral match (PSM) error rates with a 1% FDR were determined using the target-decoy strategy coupled to Percolator modelling of true and false matches.

Both unique and razor peptides were used for quantitation. Signal-to-noise (S/N) values were used to represent the reporter ion abundance with a co-isolation threshold of 50% and an average reporter S/N threshold of 10 and above required for quantitation from each MS3 spectra to be used. The summed abundance of quantified peptides were used for protein quantitation. The total peptide amount was used for the normalisation. Protein ratios were calculated from medians of summed sample abundances of replicate groups. Standard deviation was calculated from all biological replicate values. The standard deviation of all biological replicates lower than 25% was used for further analyses.

Differentially regulated proteins were identified using a linear-based model (limma) (Ritchie et al. 2015) on the normalized log2 protein abundance. P value < 0.05 threshold was used as significance threshold. The Generally Applicable Gene-set Enrichment (GAGE) was used to retrieve the enriched processes (Luo et al. 2009). Several databases from MSigDB were used including Hallmark, Reactome, GO and immunologic signatures gene-sets. P-value <0.05 was used as significance threshold (Liberzon et al. 2015).

### Statistical analysis

All data were tested for normality applying the D’Agostino and Pearson or Shapiro-Wilk test. For the comparison of two groups, an unpaired two-tailed Student’s t test was applied. For data not meeting the criteria for normality, the Mann-Whitney or Wilcoxon signed-rank test was applied. For the analysis of more than two groups, one-way analysis of variance (ANOVA) was applied in case of normally distributed data. Correcting for multiple comparison was performed by Dunnett’s test (comparing all groups to one control group). Nonparametric data were analyzed using the Kruskal-Wallis test followed by Dunn’s test (comparing all groups to one control group) to correct for multiple comparisons. Grouped analyses were tested using two-way ANOVA. To correct for multiple comparisons, Dunnett’s (comparing groups to respective control group inside of one row), Sidak’s (comparing groups to respective control group inside of one column or comparing two groups in one row) or Tukey’s (comparing all groups to each other) test was applied. If data were analyzed compared to a hypothetical value of 1 or 100, we used the one-sample t-test for normally distributed data and the one-sample Wilcoxon test when nonparametric testing was suggested. Log-rank test (Mantel-Cox) was used to calculate significance of differences between survival curves. Statistical analysis was performed using GraphPad Prism (v9, Graph-Pad Software). Applied analyses and statistical significances are indicated in the corresponding figures and figure legends. All data are represented as means  ±  SEM. Differences with p≤0.05 were considered statistically significant. ns=non significant, * p≤0.05, ** p≤0.01, *** p≤0.001, **** p≤0.0001. All data values and the corresponding statistical tests of each graph are available.

## Acknowledgements

We acknowledge the excellent scientific and technical assistance of the Signalling Factory Core Facility staff and the Life Imaging Center (LIC) in the Center for Biological Systems Analysis (ZBSA), both of the University of Freiburg, E. Donnadieu for teaching us to perform migration experiments on viable tumor slices and M. Hofmann for providing the α-PD-1 antibody Nivolumab. We thank G. Siegers, C. Normann and G. Fiala for knowledge transfer and technical support. This study was supported by the German Research Foundation (DFG) through BIOSS - EXC294 and CIBSS - EXC 2189 to S.M.; SFB850 (C10 to S.M.), SFB1479 (Project ID: 441891347 - P15 to S.M.), SFB1160 (Project B01 to SM), and FOR2799 (MI1942/3-1 to SM). K.R. was supported by DFG through GSC-4 (Spemann Graduate School), MB was supported by SFB850 subprojects C9 and Z1, SFB1479 (Project ID: 441891347 - S1), SFB1160 (Project Z02), SFB1453 (Project ID 431984000 - S1) and TRR167 (Project Z01), the German Federal Ministry of Education and Research by MIRACUM within the Medical Informatics Funding Scheme (FKZ 01ZZ1801B).

## Author contributions

K.R. and S.M. designed the study. K.R. planned and performed most experiments. J.M. performed the BCSC xenograft transplantation in mice. J.S. participated in the *in vivo* experiments, the BCSC culture and in the experiments with xenograft-derived cells. K.M.K. and M.Z. helped with viral particle production. O.T. performed the analysis of γδ T cell migration experiments. M.S. and H.Z. performed the proteomic study. G.A. and M.B. bioinformatically analyzed the proteomic data. J.M., W.W. and M.B. provided intellectual input and critically read the manuscript. K.R. and S.M. wrote the manuscript and S.M. supervised the study. All authors discussed the results and conclusions drawn from the studies.

## Competing Interests statement

The authors declare no competing interests.

---

**Extended Data Fig. 1:**
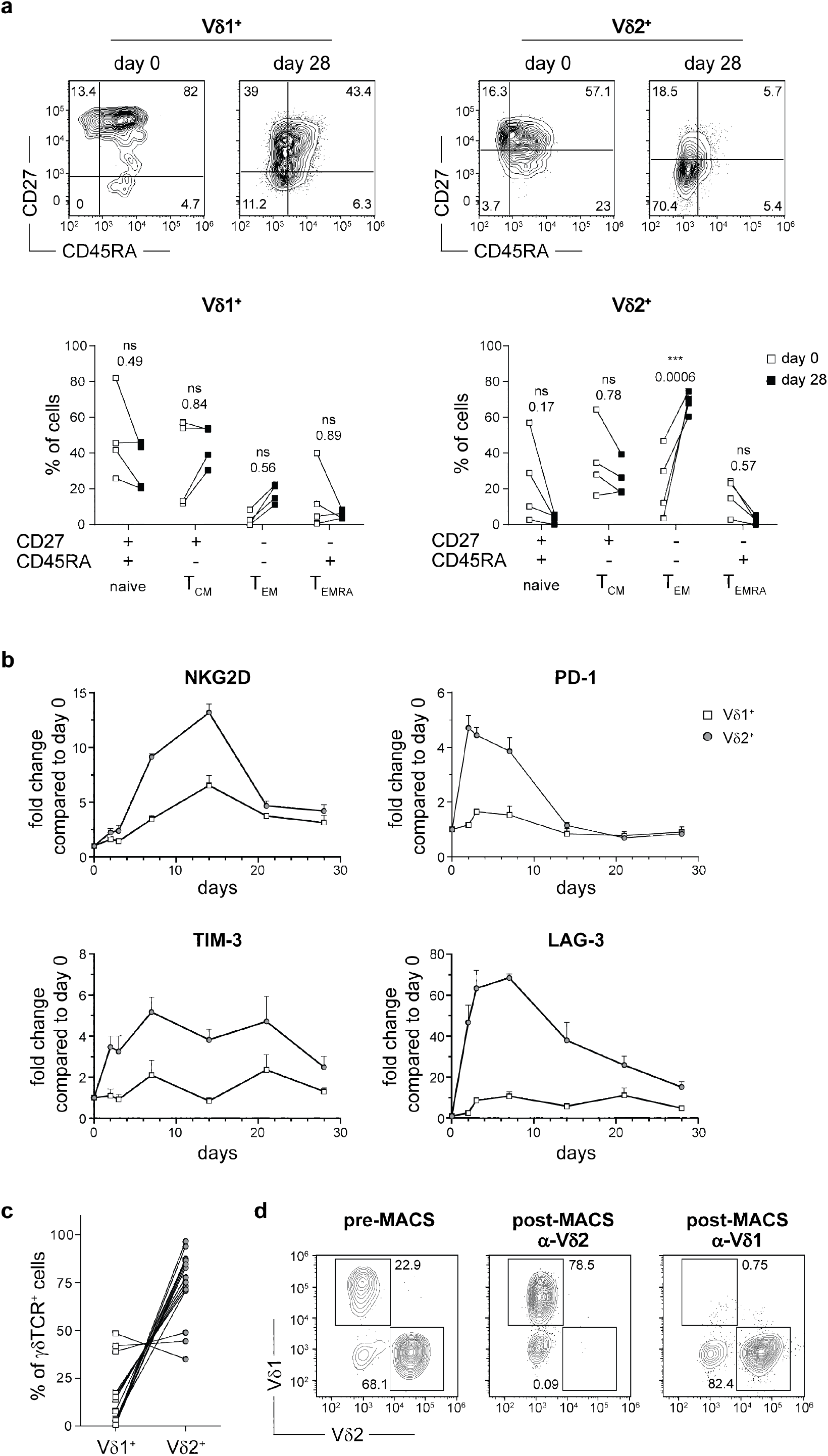
Expanded human γδ T cells are mainly of the central memory and effector memory phenotype. **(a)** Flow cytometry-based analysis of Vδ1^+^ and Vδ2^+^ T cells on day 0 and day 28 after peripheral blood mononuclear cell (PBMC) stimulation with 10 µg/ml Concanavalin A (ConA), 10 ng/ml IL-2 and 10 ng/ml IL-4. Cells were stained and gated to distinguish Vδ1^+^ and Vδ2^+^ T cells and phenotyped for the expression of CD27 and CD45RA: naïve T cell, CD27^+^CD45RA^+^; central memory T cell (T_CM_), CD27^+^CD45RA^-^; effector memory T cell (T_EM_), CD27^-^CD45RA^-^; CD45RA^+^ effector memory T cell (T_EMRA_), CD27^-^CD45RA^+^. Gates were based on fluorescence-minus-one (fmo) controls. Shown are representative dot plots from one healthy donor (upper panels) and statistical analysis of pooled results of four healthy donors stained simultaneously (lower panels). Two-way ANOVA followed by Sidak’s post hoc test comparing stimulated or co-cultured cells to the corresponding medium control. **(b)** PBMCs stimulated as in (a) were stained and gated to distinguish Vδ1^+^ and Vδ2^+^ T cells and the expression of NKG2D, PD-1, TIM-3 and LAG-3 was monitored over time. Data were normalized to the mean fluorescence intensity (MFI) at day 0. Pooled data of three healthy donors from one experiment are shown (means ± SEM). **(c)** Analysis of the percentage of Vδ1^+^and Vδ2^+^ T cells among γδTCR^+^ gated cells. Shown are the results for 15 independent expansion cultures from different donors. **(d)** Representative flow cytometry dot plots of γδ T cell cultures before and after magnetic activated cell sorting (MACS). Vδ2^+^ or Vδ1^+^T cells were depleted from cultures by using α-Vδ2 or α-Vδ1 antibodies, respectively. Shown are γδTCR^+^ gated cells. *** p≤0.001.

**Extended Data Fig. 2:**
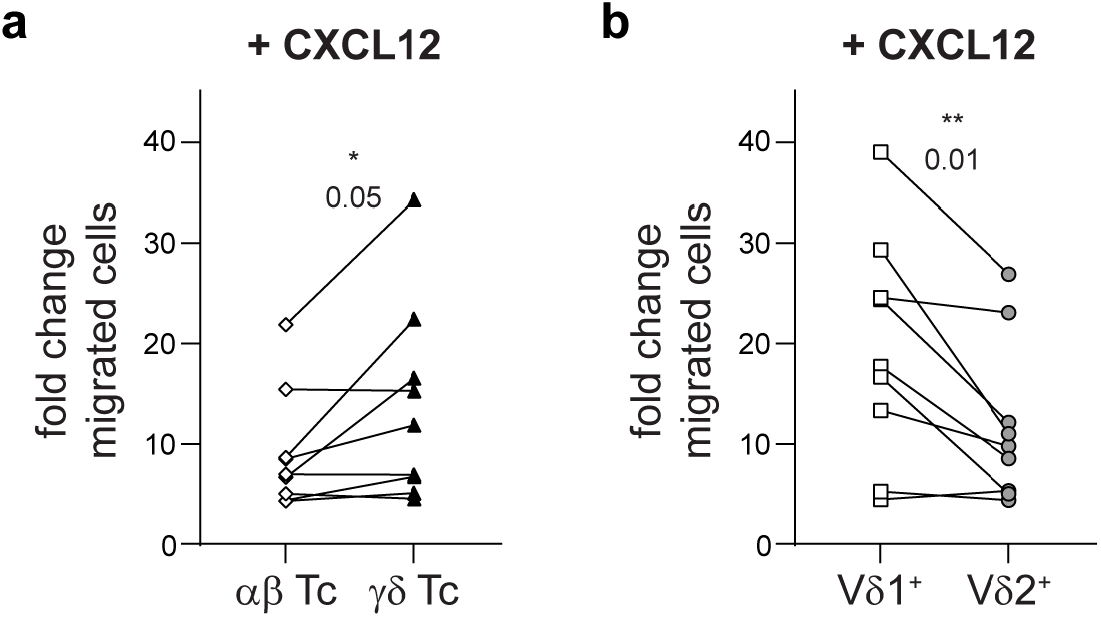
T cell migration towards the chemokine CXCL12. **(a)** Migration of αβ T cells (CD3^+^γδTCR^-^) and γδ T cells (CD3^+^γδTCR^+^) in response to CXCL12 (50 ng/ml) was determined in a transwell assay as described in Fig. 2 (a). Basal migration towards medium was set to 1.0 and fold changes from three independent experiments using the same three healthy donors in each experiment were pooled. Wilcoxon signed-rank test. **(b)** Migration of Vδ1^+^ (CD3^+^γδTCR^+^Vδ1^+^) and Vδ2^+^ (CD3^+^γδTCR^+^Vδ2^+^) T cells in response to CXCL12 (50 ng/ml) was determined in a transwell assay, which was analyzed as detailed in (a). Wilcoxon signed-rank test.* p≤0.05, ** p≤0.01.

**Extended Data Fig. 3:**
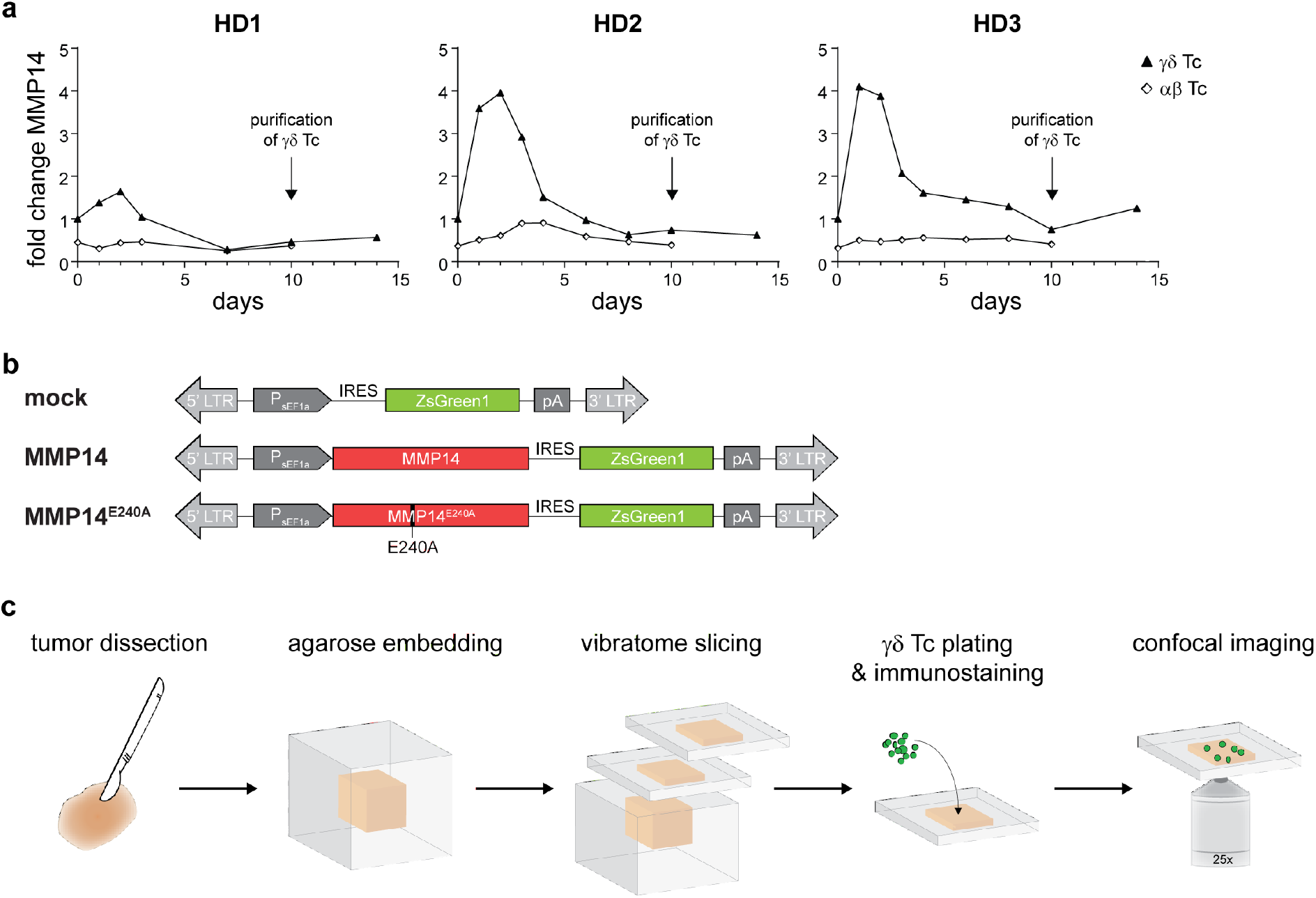
MMP14 expression inγδ T cells. **(a)** Flow cytometry-based analysis of endogenous MMP14 expression in γδ (CD3^+^γδTCR^+^) and αβ (CD3^+^γδTCR^-^) T cells after the stimulation of PBMCs with ConA (10 µg/ml), IL-2 (10 ng/ml) and IL-4 (10 ng/ml). The MFI of MMP14 expression on γδ T cells on day 0 was set to 1 for normalization. αβ T cells were depleted via MACS from the cultures on day 10 after initial stimulation (arrow). Results for three healthy donors of γδ T cells are shown. (**b**) Schematic diagram of the lentiviral vectors used to express MMP14 or the catalytically inactive mutant MMP14^E240A^. **(c)** Experimental setup to analyze γδ T cell migration in viable slices of BCSC5 xenograft tumors. BCSC5 xenograft tumors were embedded in agarose and viable tissue slices of 350 µm thickness were cut using a vibratome. Slices were stained with fluorophore-conjugated antibodies and CMFDA-labeled γδ T cells were plated on top of the tumor slices. After allowing γδ T cells to infiltrate the tissue for 30 min, infiltrated cells were imaged by confocal microscopy. Modified from previously published protocols (Salmon et al. 2012; Salmon et al. 2011). CMFDA, 5-Chloromethylfluorescein diacetate; IRES, internal ribosomal entry site; LTR, long terminal repeat; pA, polyA; sEF1α, short elongation factor 1-alpha.

**Extended Data Fig. 4:**
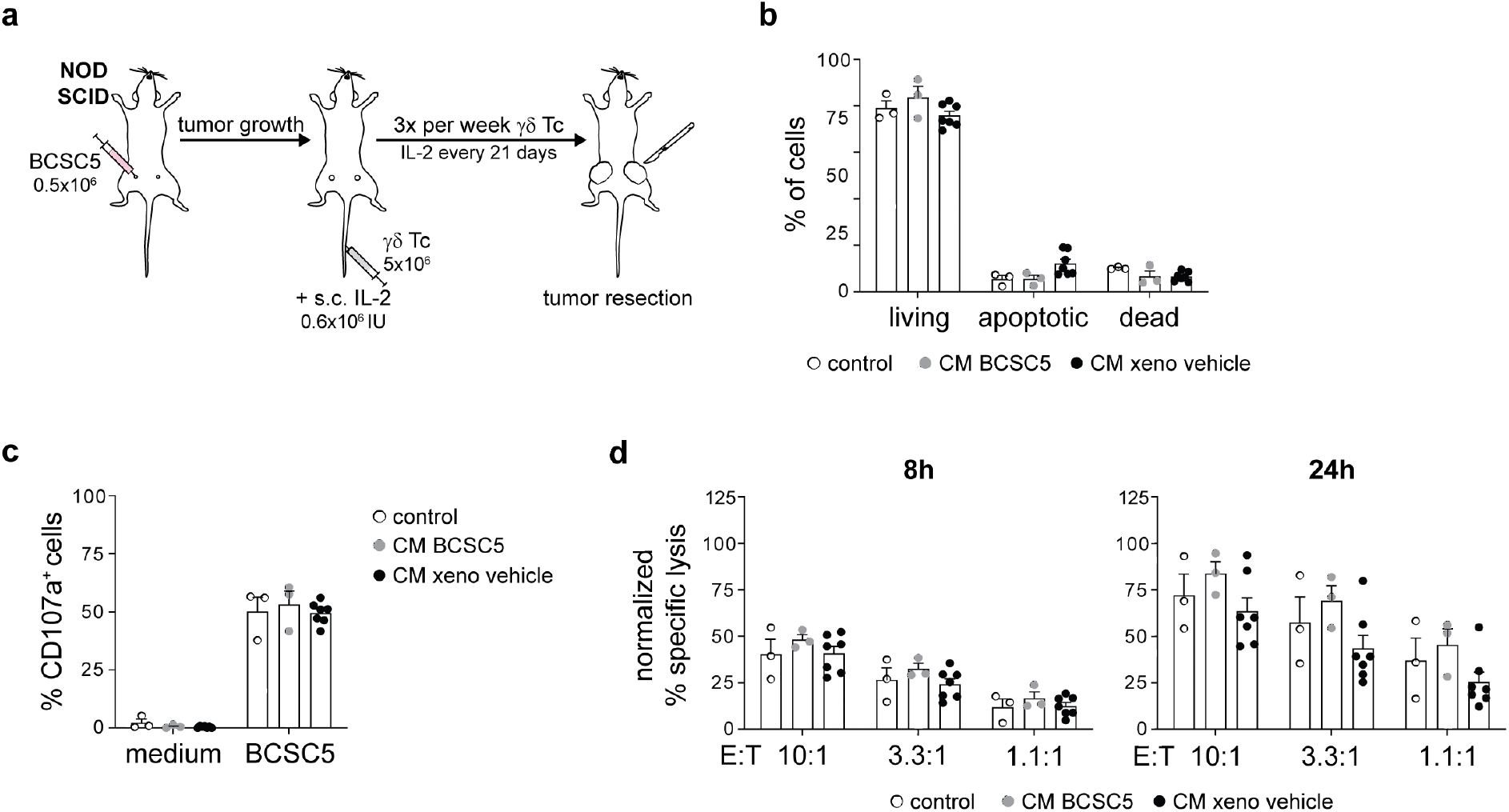
γδ T cell therapy of BCSC5 xenografts in NOD SCID mice. **(a)** Schematic illustration of BCSC5 transplantation and γδ T cell immunotherapy in immunocompromised mice. 0.5x10^6^ BCSC5 cells were orthotopically transplanted into each fat pad of the two #4 mammary glands of 6-8 weeks old female mice. Treatment start was defined for each mouse individually when the first tumor reached a volume of at least 8 mm^3^. 5x10^6^γδ T cells were injected intravenously three times per week. In addition, mice received 0.6x10^6^ IU IL-2 (Proleukin S) on the day of treatment start and every 21 days until the end of the experiment. The end of the experiment was defined by a tumor volume of 800 mm^3^. (**b**) Flow cytometry-based assessment of γδ T cell viability after exposure to conditioned medium (CM) from BCSC5 culture cells or xenograft-derived tumor cells (xeno). γδ T cells were cultured in CM for 24 h. CM was removed and cell viability was analyzed 24 h later. Living: Annexin V^-^/propidium iodide (PI)^-^, apoptotic: Annexin V^+^/PI^-^, dead: Annexin V^+^/PI^+^. Results for two healthy donors of γδ T cells from three independent experiments were pooled (means ± SEM). Two-way ANOVA followed by Dunnet’s post hoc test comparing CM samples to respective controls. Differences were not statistically significant. **(c)** Flow cytometry-based analysis of degranulation by γδ T cells in response to BCSC contact for 3 h. γδ T cells were cultured in CM from BCSC5 culture cells or xenograft-derived tumor cells for 24 h and then used for degranulation assays. The percentages of CD107a^+^ cells of γδTCR^+^-gated cells for two healthy donors of γδ T cells from three independent experiments were pooled (means ± SEM). Two-way ANOVA followed by Dunnet’s post hoc test comparing CM samples to respective controls. No significant differences were obtained. **(d)** *In vitro* killing of luciferase-expressing BCSC5 by γδ T cells after 8 h and 24 h at different effector to target (E:T) ratios. γδ T cells were cultured in CMfrom BCSC5 culture cells or xenograft-derived tumor cells for 24 h and then used for cytotoxicity assays. Results for two healthy donors of γδ T cells from three independent experiments were pooled (means ± SEM). Two-way ANOVA followed by Dunnet’s post hoc test comparing CM samples to respective controls. No significant differences were obtained.

**Extended Data Fig. 5.**
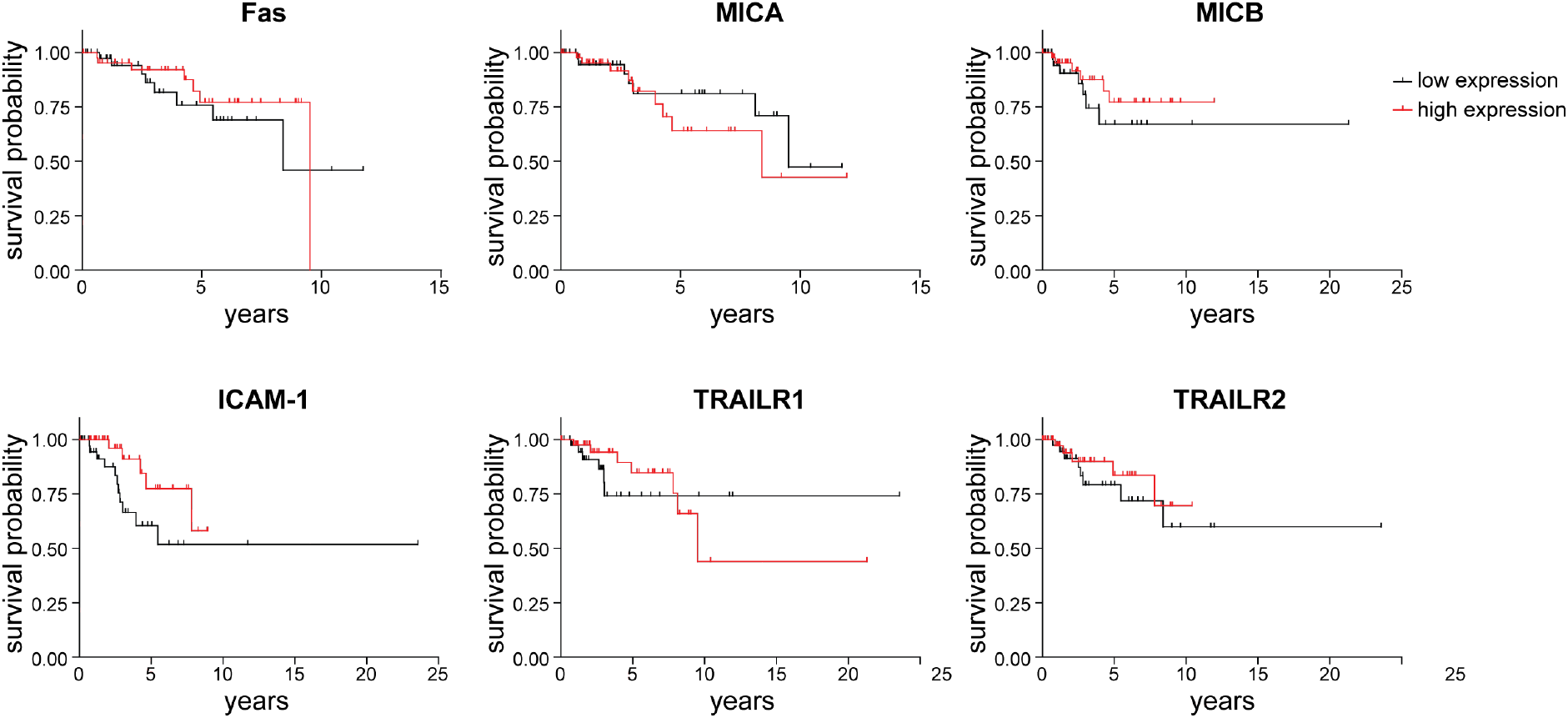
(corresponds to Fig. 6): Killing of BCSC5 by γδ T cells requires multiple ligand-receptor interactions. **(a)** Cox regressions of progression-free survival for TNBC patients sorted by high (upper-quartile) and low (lower-quartile) expression of the indicated proteins. All individual proteins but MICA showed negative β cox coefficient indicating that patients with high expression exhibit a better survival prognosis. However, none of the individual cox p value was statistically significant (<0.05).

**Extended Data Fig. 6.**
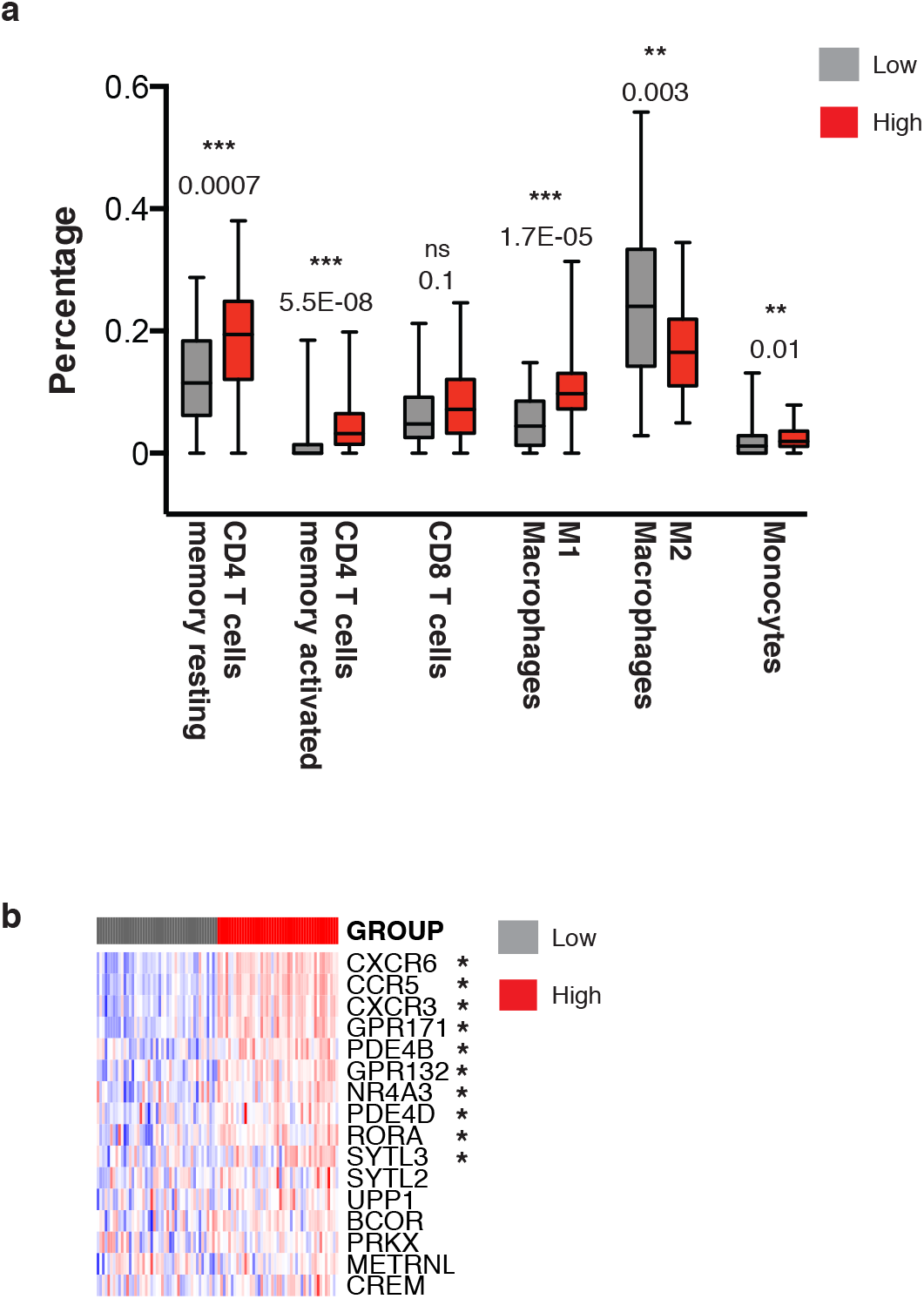
(corresponds to Fig. 7): Immune profiling of TNBC patients. **(a)** TNBC patients were sorted by high (upper-quartile) and low (lower-quartile) average clustered expression of BTN2A1, BTN3A1, Fas, MICB and ICAM-1 and the deep deconvolution CIBERSORT algorithm to deduce the immune cell composition. * p≤0.05, ** p≤0.01, *** p≤0.001. **(b)** Row wise Z-score heatmap for a gene set specific for human γδ T cells according to (Tosolini et al. 2017). Out of 23 genes forming this set, only 16 were annotated in the database. Significantly different expressed genes are indicated with *.

## Notes

### Competing Interest Statement

The authors have declared no competing interest.

